# A single-cell atlas of *Plasmodium falciparum* transmission through the mosquito

**DOI:** 10.1101/2020.10.11.333179

**Authors:** Eliana Real, Virginia M. Howick, Farah Dahalan, Kathrin Witmer, Juliana Cudini, Clare Andradi-Brown, Joshua Blight, Mira S. Davidson, Sunil Kumar Dogga, Adam J. Reid, Jake Baum, Mara K. N. Lawniczak

## Abstract

Malaria parasites have a complex life cycle featuring diverse developmental strategies, each uniquely adapted to navigate specific host environments. Here we use single-cell transcriptomics to illuminate gene usage across the transmission cycle of the most virulent agent of human malaria – *Plasmodium falciparum*. We reveal developmental trajectories associated with the colonisation of the mosquito midgut and salivary glands and elucidate the transcriptional signatures of each transmissible stage. Additionally, we identify both conserved and nonconserved gene usage between human and rodent parasites, which point to both essential mechanisms in malaria transmission and species-specific adaptations potentially linked to host tropism. Together, the data presented here, which are made freely available via an interactive website, establish the most complete atlas of the *P. falciparum* transcriptional journey to date.

**One sentence summary:** Single-cell transcriptomics of *P. falciparum* transmission stages highlights developmental trajectories and gene usage.

---

Malaria parasite transmission to and from the mosquito vector is accompanied by a precipitous drop in parasite population numbers (*1, 2*). The bottleneck that transmission affords has, therefore, long been considered an attractive target for antimalarial or vaccine interventions. In a highly orchestrated series of developmental transitions taking place inside the mosquito, parasites taken up during a blood meal complete sexual development, fertilisation, and recombination; differentiate into invasive ookinetes that colonise the midgut; develop into oocysts where population expansion occurs; and emerge as invasive sporozoites that fill the mosquito salivary glands and can be transmitted to a new host during a mosquito bite (*3*). After a relatively brief stint in the skin, some injected sporozoites will reach the liver, where they undergo a phase of profound population expansion, followed by a release of merozoites that enter the bloodstream (*4*). This heralds the beginning of the cyclical asexual cycle, where merozoites invade erythrocytes and develop intra-erythrocytically, further expanding parasite numbers, with some parasites also developing into sexual, transmissible forms that can perpetuate the transmission cycle when taken up in the bloodmeal of a mosquito.

Single-cell RNA sequencing (scRNA-seq) has transformed our ability to resolve cell-level heterogeneity in complex cell populations. Recent scRNA-seq views across the full life cycle of *Plasmodium* parasites, the etiological agents of malaria, have captured fine scale developmental transitions driving progression through the life cycle at unprecedented resolution (*5–9*). The Malaria Cell Atlas (*6, 8*) was established as a data resource and website to provide an accessible route for scientists to explore patterns of gene expression in individual parasites across multiple developmental stages and parasite species. However, its application to *P. falciparum*, the species responsible for the vast majority of human death and disease has been thus far limited to the blood stages. Here we add a new chapter to the atlas, completing a scRNA-seq survey of the transmission stages of *P. falciparum* from the sexual forms transmitted to the *Anopheles* mosquito to the invasive sporozoite delivered to the human host. Understanding these key developmental transitions in parasite biology is essential in order to devise novel strategies for tackling malaria, a task rendered more urgent by the ongoing spread of antimalarial drug resistance and lack of a vaccine that affords long term protection.

To capture human malaria transmission at single-cell resolution, we assembled a comprehensive scRNA-seq data set across key stages of the *P. falciparum* life cycle (Fig. 1A). Distinct morphological forms of the parasite were purified based either on the expression of known stagespecific markers (circumsporozoite protein (CSP) and ookinete surface protein P25 (Pfs25) for sporozoites and ookinetes, respectively) or the RNA and DNA content (stage V gametocytes) (Fig. S1–3). After excluding poor quality cells through stringent quality control steps (Fig. S4, Table S1), our data set comprised 1467 single cell transcriptomes spanning the broad developmental transitions that underpin host-to-vector and vector-to-host transmission. We next integrated this transmission data set with our published collection of 161 single-cell transcriptomes of mixed *P. falciparum* asexual stages (*6*), which enabled us to reconstitute the *P. falciparum* life cycle almost entirely, with the exception of exo-erythrocytic schizogony (liver stages). In order to visualise *P. falciparum* single cells across the life cycle, we performed dimensionality reduction using UMAP and clustered cell transcriptomes using Louvain clustering (*10–12*). As expected, cells from the same stage grouped together in two-dimensional space (Fig. 1B) and formed distinct cell clusters (Fig. 1C). We were also able to observe clear cellular trajectories that are likely to reflect within-stage developmental paths (Fig. 1B, C). For example, the distribution of asexual parasites in twodimensional space clearly shows developmental progression along the IDC (intraerythrocytic developmental cycle) (fig. S5) (*6*). Likewise, the trajectory displayed by ookinete cells is consistent with a range of developmentally intermediate states along the known ookinete maturation path (fig. S6) (*13*).

**Figure 1.**
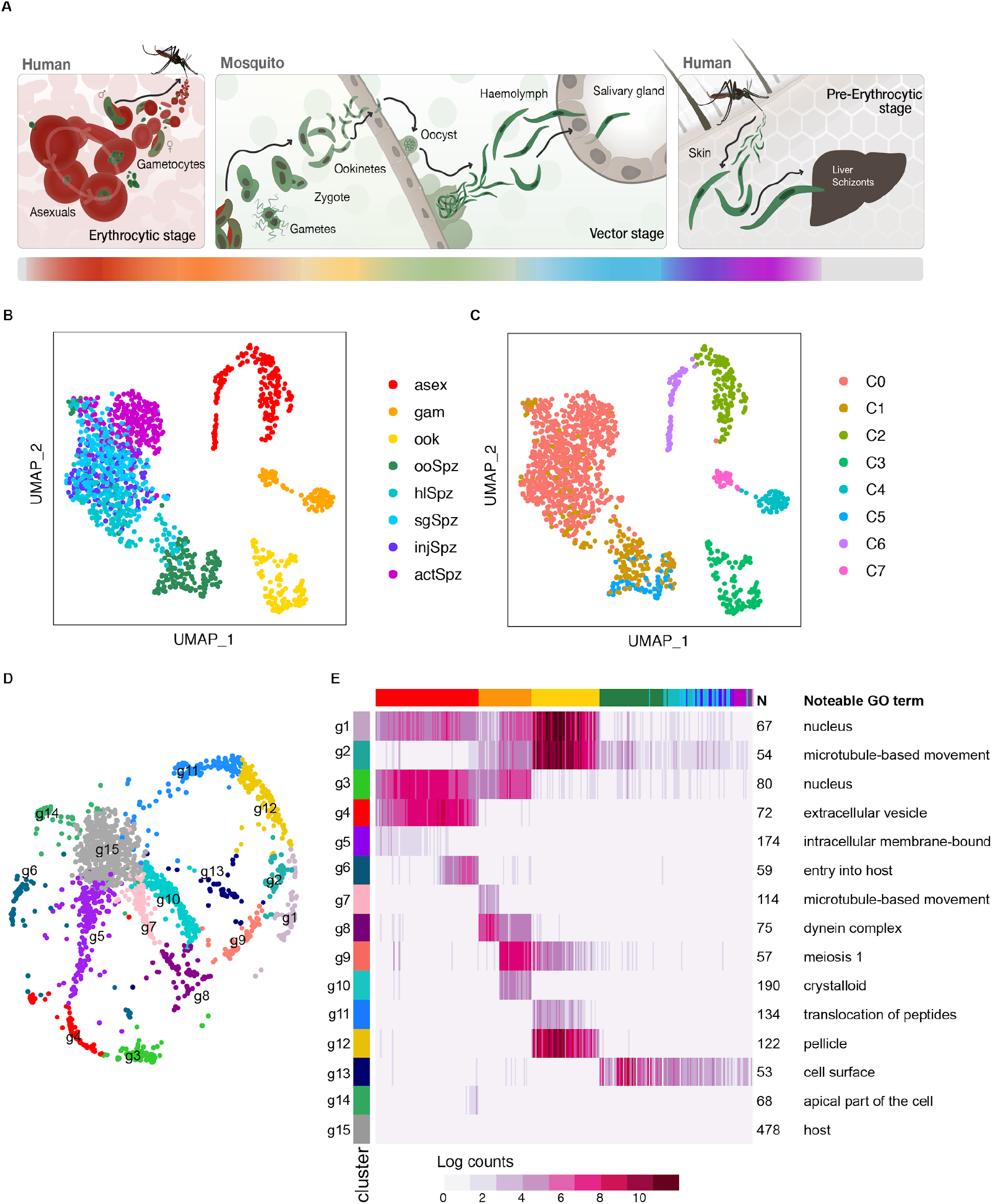
A single-cell transcriptome atlas of *P. falciparum* transmission. **(A)** The life cycle of *P. falciparum* includes asexual replication and sexual development in red blood cells (erythrocytic stages). Sexual stage gametocytes are then taken up in the mosquito blood meal, where they undergo fertilization and develop into the ookinete form that crosses the mosquito midgut epithelium and begins sporogonic development along the basal lamina as an oocyst. Sporozoites are released into the hemolymph from mature oocysts where they travel to and invade the salivary glands. A subset of these sporozoites may be injected into the next human host where they migrate through the dermis on their way to the liver (pre-erythrocytic stages). The colour bar below represents the different life cycle stages reflected in **B**. **(B)** UMAP of single-cell transcriptomes from across asexual and transmission stages of the parasite. Stage abbreviations: asex (asexual blood stages), gam (male and female gametocytes), ook (ookinete), ooSpz (oocyst sporozoites), hlSpz (hemolymph sporozoites), sgSpz (salivary gland sporozoites), injSpz (sporozoites released by mosquito bite), actSpz (activated sporozoites). Asexual stage transcriptomes have been previously published (*6*). **(C)** Louvain clustering of single-cell transcriptomes identified 8 distinct clusters. In order to get a more equal distribution of numbers of cells across clusters, clusters C0 and C1 were downsampled to 100 cells each prior to making the gene graph **(D, E)**. **(D)** A k-nearest neighbors (kNN) force-directed graph of 1797 highly variable genes across cell types. Each node represents a gene. Nodes are clustered by their graph-based spectral clustering assignment. **(E)** A heatmap of mean expression of the genes in each cluster across 623 subsampled cells. Cells are ordered by their developmental progression along the life cycle. Cells from asexual, ookinete and sporozoite stages were ordered according to their computed pseudotime values (Fig. S5, S6). Gametocytes are grouped based on cluster assignment from C, where C7 are male gametocytes and C4 are female/early gametocytes (see Fig. S2 for more detailed analysis). The male gametocyte cells are ordered before the female gametocytes in the heatmap. Stages are represented by the colours in A, B.

To gain further insight into the gene expression patterns that define each cellular state, 1797 highly variable genes in the data set, determined using a dropout-based feature selection method in scmap (*14*), were assigned to one of 15 gene co-expression modules using graph spectral clustering (denoted with g, Fig. 1D). Most gene clusters showed stage-specific expression patterns and were enriched for GO terms expected for each stage (Fig. 1E). Other gene clusters were more ubiquitously expressed, possibly reflecting shared functional strategies across the life cycle. For instance, cluster g1, which features two putative readers of post-translational histone modifications, namely bromodomain protein 1 (BDP1) and the EELM2-containing PF3D7_0519800, contains genes that tend to be expressed throughout the IDC and sexual stages, where epigenetic control of developmental processes has long been recognised (*15, 16*). Here we find that these genes are also expressed in ookinetes, implying that epigenetic mechanisms may be more widely used across the life cycle (*17*). Both BDP1 and EELM2 proteins have known or predicted links to apicomplexan AP2 transcription factors (ApiAP2) (*18–20*). Significantly, two ApiAP2 TFs for which interactions with protein complexes containing BDP1 or EELM2 have been experimentally demonstrated are also expressed in this cluster (PfAP2Tel/SP3 and PF3D7_1107800, respectively) (*19, 20*). Additionally, BDP1 has been shown to associate with the promoter regions of seven ApiAP2 TFs (*18*) whose expression we find to be correlated with gene clusters associated with asexual development and host-to-vector transmission (Fig. S7), thus expanding the reach of this gene regulatory network. Several other genes without predicted functions are co-transcribed in cluster g1 and as such could represent additional links between chromatin remodelling/targeting and AP2-dependent gene expression.

The majority of genes in the *P. falciparum* genome have one-to-one orthologs with *P. berghei* (80%). We therefore explored whether any clusters were enriched for genes that lack orthologs between the two species. This analysis revealed that every cluster had some clade-specific genes present, possibly reflecting species-specific adaptive mechanisms present across the life cycle (Fig. S8A, B, Fig. S9). Cluster g15, which is the largest gene cluster by more than two-fold showed low levels of expression across the life cycle and had the highest proportion of genes lacking orthologs, with a large number of *P. falciparum* variant multigene families (*e.g*. stevors, rifs) associated with host interactions in blood stages (GO:0018995, Bonferroni adjusted *p*-value = 3.58 x 10^−9^) (*21*). Clusters associated with mosquito stages and vector-to-host transmission (g1, g2, g9, g12, g13), in contrast, had proportionally more genes with *P. berghei* orthologs. Despite being enriched for one-to-one orthologs, two of these clusters (g2, g13) showed the lowest amino acid conservation scores across the gene graph (Fig. S8C, D). Both clusters feature genes encoding surface proteins with essential roles in transmission, suggesting that protein interactions that underpin transmission might have been shaped by coevolution of parasite and host and may account, at least in part, for host specificity/tropism in *Plasmodium* infections.

To further explore conservation of genes involved in *Plasmodium* transmission, we compared *P. falciparum* transmission stages with the equivalent *P. berghei* stages from Howick *et al*. using scmap (*14*). The two data sets showed a good overall correlation, with *P. falciparum* cells mapping to cells of the same stage in the *P. berghei* reference data set (Fig. S10A, B) (*8*). As expected, cells that were not represented in the *P. berghei* transmission analysis (early sporozoite developmental stages), showed lower similarity across species than those that were common to the two data sets (Fig. S10C). We next used an unsupervised integration and graph-based clustering approach in Seurat (*11, 12*) to elucidate gene expression patterns across the transmission stages of the two species. Using this approach, cells from the two data sets could be matched to four clusters, with cells mixing by stage in the life cycle (Fig. 2A, B, C). A fifth cluster (C1) was populated by *P. falciparum* midgut and hemolymph sporozoites that were matched by very few cells in the *P. berghei* data set (Fig. 2A, B, C). The global structure of the integrated data set remained largely unchanged compared to the original *P. falciparum* subset, underlining the robustness of the cell identity correlations. For each cluster, we identified marker genes for each stage that are conserved between species (Fig. 2D, E). Among genes that represent the conserved transcriptome (Fig. 2D, E, Table S1), 30 to 40% have no known function, but, as exemplified for the ookinete subset (Fig. S6F), we can begin to make predictions about gene function by surveying correlated gene expression patterns along highly resolved developmental trajectories. The same strategy could help to fast-track discovery of new candidates for therapeutic development, for example, by uncovering genes with expression profiles that are highly correlated with those of validated vaccine targets (e.g. Pfs25, CSP) (*22*).

**Figure 2.**
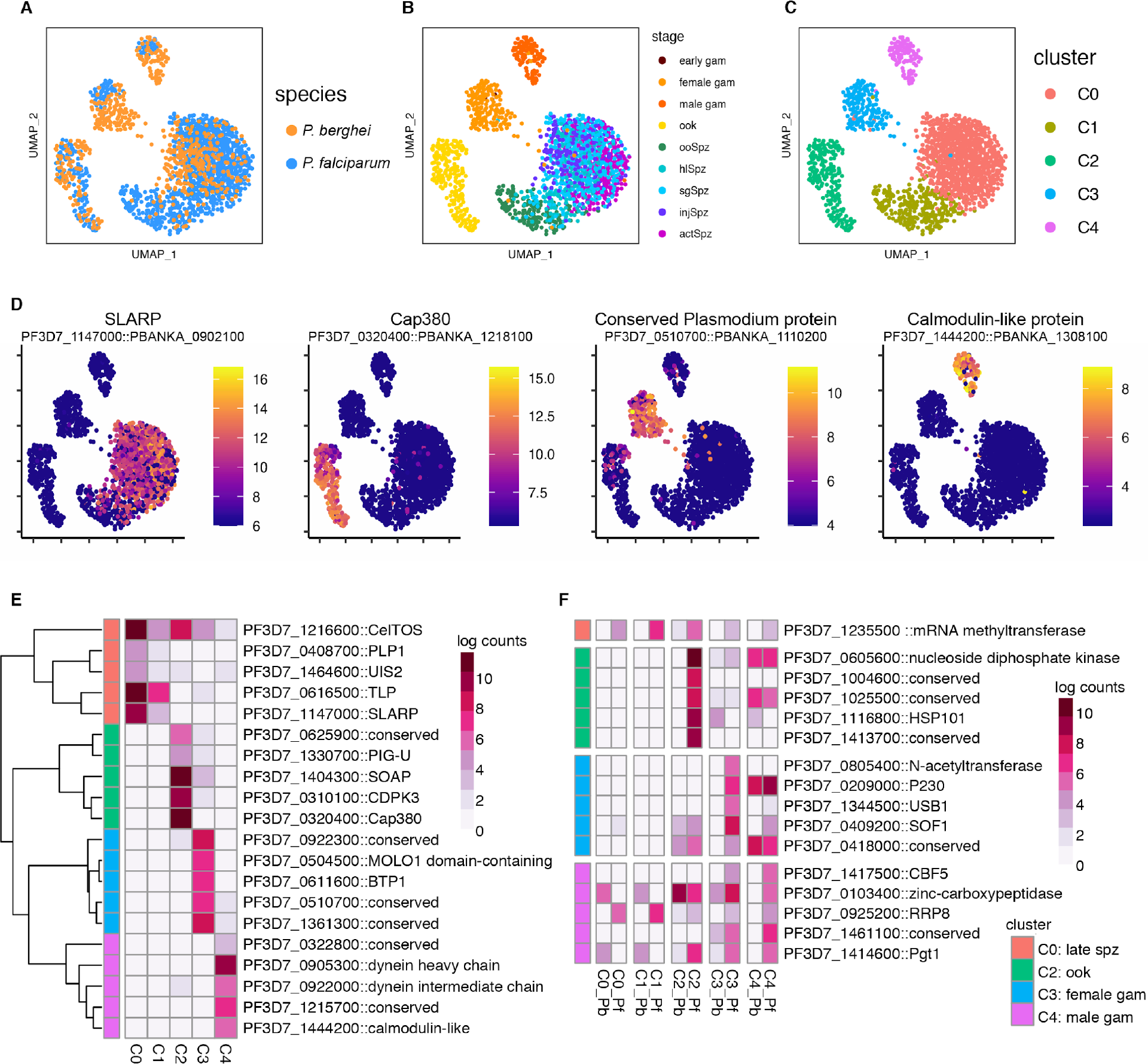
Integration of *P. falciparum* and *P. berghei* transmission datasets. To identify conserved and divergent patterns of gene expression *P. falciparum* single-cell data was integrated with the *P. berghei* transmission stage data using one-to-one orthologs. **(A-C)** UMAP of integrated data colored by species **(A)**, collection stage **(B)**, and cluster assignment **(C)**. **(D)** Expression (log counts) of the top conserved marker gene, based on the lowest maximum p-value across the two species, for clusters 0 (SLARP), 2 (Cap380), 3 (conserved *Plasmodium* protein PF3D7_0510700), and 4 (calmodulin-like protein) on the UMAP. **(E)** Heatmap of the top five conserved marker genes, based on the lowest maximum p-value across the two species, for each cluster. The expression values represent the mean log counts for each cluster. **(F)** A heatmap of the top one-to-one orthologs that are more highly expressed in *P. falciparum (*Pfa) compared to *P. berghei (*Pbe) (based on adjusted p-value and expression in at least 70% of *P. falciparum* cells). The expression values represent the mean log counts for each species in each cluster. Heatmaps are annotated with the *P. falciparum* gene identifier for the ortholog and a manual annotation. Genes annotated as conserved are conserved proteins with unknown function in *Plasmodium*.

In addition to shared patterns in genome activity with broadly conserved *Plasmodium* genes, our integrated analysis also revealed non-conserved patterns of gene expression. In all stages we identified genes whose transcripts could be detected in over 70% of *P. falciparum* cells but showed no or only residual expression in *P. berghei* cells of the same stage (Fig. 2F). Together with the genes that do not have orthologs in *P. berghei* (Fig. S8A, B), these differentially expressed genes may represent species-specific adaptations that allow *P. falciparum* to interact optimally with the diverse host environments it meets as it transmits from vector to host and vice-versa. For example, *P. falciparum* ookinetes (C2_Pf in Fig. 2F) displayed the most striking pattern of species specific expression with 26 genes (several of which have no known function) expressed in greater than 70% of *Pf* cells and in less than 10% of *Pb* cells (File S1). Conversely, in sporozoites, a single one-to-one ortholog – encoding a putative mRNA methyltransferase (PfMT-A70.2) – showed *P. falciparum-specific* expression (Fig. 2F). PfMT-A70.2 was recently identified as part of a methyltransferase complex that modulates gene expression during the IDC through N^6^-methyladenosine methylation of mRNA transcripts (*23*). Our observation that PfMT-A70.2 expression is highly correlated with the mosquito stages in *P. falciparum* but not in *P. berghei* (Fig. 2F), hints at unexplored species-specific mechanisms to fine-tune gene expression during the parasite journey in the mosquito.

Understanding how sporozoite infectivity towards the human host develops is likely to inform the identification of new antimalarial and vaccine candidates. We therefore sampled these stages more densely to capture a finely resolved developmental trajectory that recapitulated the sporozoite journey through the mosquito, with its different anatomical, cellular, and physicochemical environments (*24*). We re-clustered and oriented sporozoites along a pseudotemporal trajectory where the relative ordering of each individual cell was defined by its transcriptional state (Fig. 3A, B). This enabled us to explore fine transcriptional changes associated with a continuum of development instead of discrete time points, which were observed to be heterogeneous (Fig. 3C). The pseudotime trajectory aligned with the expected direction of development, with cells broadly segregating by their anatomic site and developmental time point (Fig. 3B, C). Differential expression (DE) analysis uncovered 301 genes with expression significantly associated with pseudotime (q-value < 0.0001). Approximately 20% of these overlapped with genes previously identified as <u>u</u>pregulated in <u>o</u>ocyst <u>s</u>porozoites (UOS) or <u>u</u>pregulated in <u>i</u>nfectious <u>s</u>porozoites (UIS) (Fig. S11A, File S1) (*25*), but many others have never been linked with sporozoite biology.

**Figure 3.**
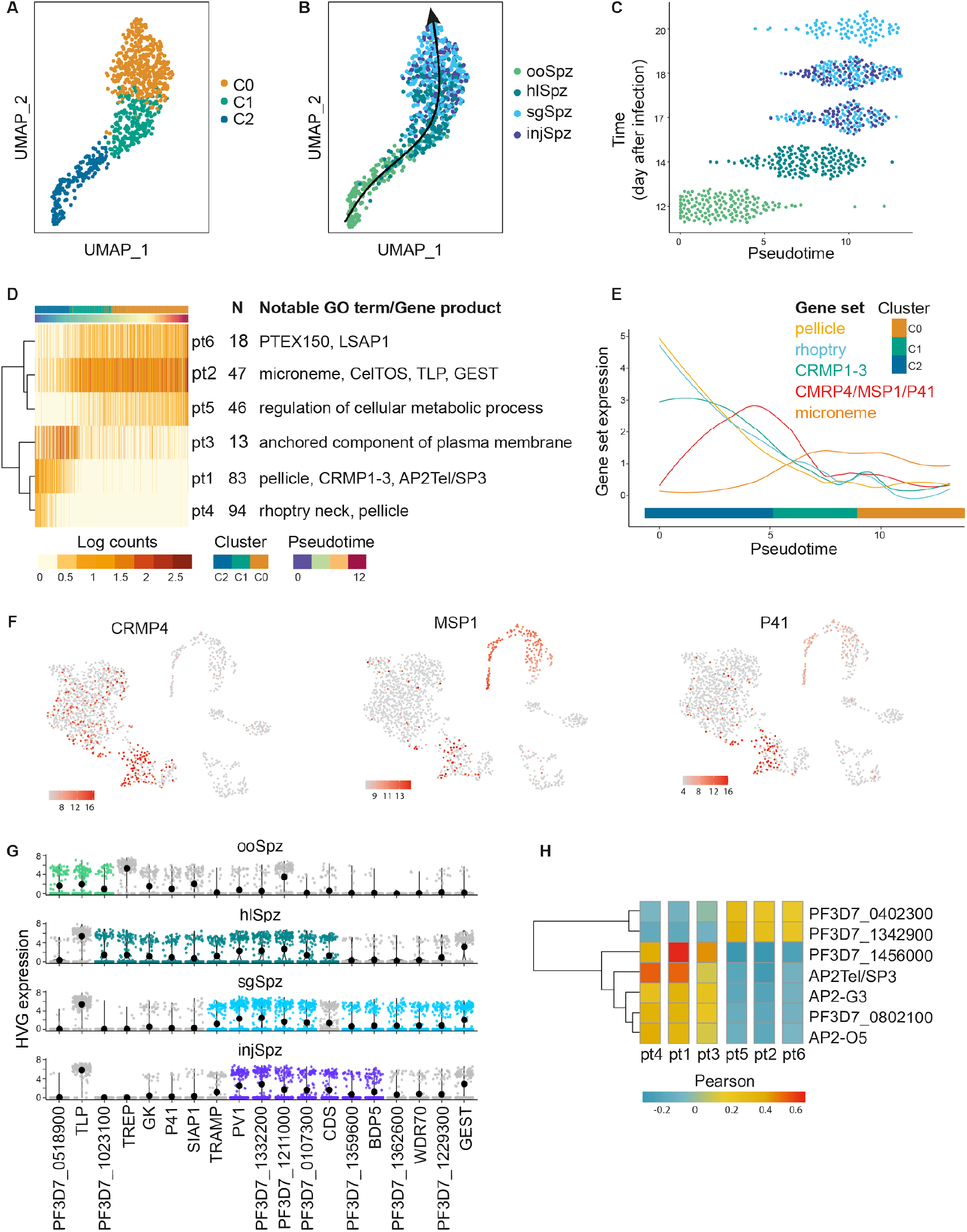
The sporozoite journey in the mosquito. **(A-B)** UMAP representation of single-cell sporozoite transcriptomes at different stages of the sporozoite developmental journey coloured by cluster assignment **(A)** and sorted stage **(B)**. The arrow in B represents the direction of development. **(C)** Sporozoites aligned along pseudotime. Overall, the pseudotime trajectory correlates with the direction of development indicated by the time of sample collection in the y axis. **(D)** Heatmap showing the mean expression of differentially expressed (DE) genes over pseudotime, arranged into six pseudotime clusters (pt1 – pt6). Each row in the heatmap represents the average expression of all genes in one cluster (the number of genes in each cluster, N, is indicated). Genes in each cluster show similar expression dynamics along the developmental trajectory. **(E)** Gene modules representative of enriched GO terms (Bonferroni adjusted *p*-value < 0.05) in pseudotime clusters are shown. Transient expression of CRMP4, p41, and MSP1 (GO term *anchored component of plasma membrane* in **D**) correlates with the release of sporozoites into the hemocoel. **(F)** UMAP representations of the complete life cycle (as shown in 1B) with the expression (log counts) of CRMP4, p41, and MSP1 highlighted. **(G)** Expression of highly variable genes (HVG) across sporozoite developmental stages. HVGs were independently determined for each stage using M3Drop with a false discovery rate < 0.05. Only HVGs that are also associated with the pseudotime trajectory are shown. HVGs are coloured by their stage association, as per B. **(H)** Heatmap of Pearson’s correlations between ApiAP2 TFs expression and pseudotime gene clusters.

To reveal patterns of gene activity along the pseudotime trajectory, we grouped genes with similar expression dynamics into co-expression modules (Fig. 3D, pt1-6). This analysis revealed two largely mutually exclusive transcriptional programmes, with the midgut-to-hemocoel transition (C2 -> C1) marking an abrupt shift in gene expression. Gene modules (pt1,4) featuring known salivary gland invasion genes (<u>C</u>ysteine <u>R</u>epeat <u>M</u>odular <u>P</u>rotein, CRMP1-2) (*26*), components of the rhoptry neck and inner membrane complex were the first to downregulate expression. This was followed by a transient peak (pt3) showing high temporal correlation with sporozoite release into the hemocoel and preceding the transcriptional changes that characterised later development (Fig. 3D, E). Of note, one of the 13 genes that constitute the pt3 module, CRMP4, has been linked to sporozoite egress in *P. berghei* (*27*). Two other genes within this module – MSP1 and the 6-Cys family member p41 (*28, 29*) – encode merozoite surface proteins that have never been associated with mosquito stages. While the function of p41 has not been elucidated, MSP1 facilitates merozoite egress by destabilising the erythrocyte submembrane cytoskeleton (*30*). Its co-regulation with CRMP4 suggests an unexpected mechanistic link between sporozoite and merozoite egress (Fig. 3F). Given their highly correlated expression, we hypothesize that genes in this module form the core of a sporozoite egress functional network and anticipate that future studies into the role of MSP1 may uncover new parallels between events that lead up to the rupture of the oocyst wall and that of the erythrocyte membrane.

After their release into the hemocoel, sporozoites experienced a dramatic shift in gene expression. Several transcripts previously identified as UIS (*25*) begin to be expressed during this stage or, in some cases (e.g. CelTOS, TLP), even before egress (Fig. 3D). Many of these transcripts appear to be under translational repression (Fig. S11D) (*25*), implying that the protein products they encode are not needed until after transmission. A small proportion of these developmentally regulated transcripts, including a translationally repressed epigenetic reader (BDP5 (*25, 31*)), show highly variable expression between individual sporozoites residing in the mosquito salivary glands or released through a mosquito bite (Fig. 3G, Table S1). Given the strong association of these highly variable genes (HVG) with sporozoite development, it is tempting to speculate that the observed differences in expression between individual sporozoites could result in different readiness levels with respect to navigating the journey to the liver and/or establishing a new replication niche once they have reached their destination.

Consistent with the key role of ApiAP2s in driving all major developmental switches during the parasite life cycle (*32*), seven ApiAP2 transcripts were found to be dynamically regulated along the sporozoite developmental trajectory (Fig. S12, Fig. 3H). Of the five ApiAP2 transcription factors expressed during late midgut sporogony (cluster C2), only one – PfAP2Tel/SP3 (PF3D7_0622900) – had been previously implicated in sporozoite development. Gene knockout studies in *P. berghei* showed that AP2Tel/SP3 has a role in sporozoite release (*33*) and indeed, we observed that PfAP2Tel/SP3 expression preceded the transcription of egress genes (Fig. 3C). Future studies should examine whether PfAP2Tel/SP3 has a role in regulating the expression of egress related genes and to what extent recent findings mapping PfAP2Tel/SP3 to the telomeric ends of *P. falciparum* chromosomes (*19*) might be linked to this putative activity. In contrast to PfAP2Tel/SP3, AP2-G3 and AP2-O5, which have known roles in gametocytogenesis and in regulating ookinete motility, respectively, have not been previously implicated in sporozoite development (*34*). The two ApiAP2 transcription factors with similar expression dynamics to PfAP2Tel/SP3, PF3D7_0802100 and PF3D7_1456000 (Fig. S12), may have redundant functions in regulating sporozoite egress, as both bind the same CA-repeat promoter element (*35*). Expression of putative transcription factors PF3D7_0420300 and PF3D7_1342900 was, in contrast, correlated with gene expression post egress (Fig. 3H). Attempts to knock out the former in *P. falciparum* have previously proved unsuccessful, suggesting PF3D7_0420300 plays an essential role in asexual development (*36*). How ApiAP2 TFs active in multiple stages of the life cycle achieve stage-specific outcomes is not well understood, but may involve combinatorial binding between members of the ApiAP2 family (*32*). Thus, our analysis expands the constellation of ApiAP2 TFs with putative roles in sporozoite development and suggests that a far more complex regulatory network than previously thought – involving specific combinations of ApiAP2 TFs possibly acting cooperatively – might be required to orchestrate the dynamic transcriptional patterns associated with the sporozoite journey in the mosquito.

As sporozoites migrate away from the site of injection, they can spend up to two hours probing their environment, actively migrating in and out of dermal fibroblasts and phagocytic cells in search of a blood vessel through which they can gain access to the bloodstream (*37–39*). Being devoid of a protective vacuole, sporozoites migrating in the dermis are particularly exposed to the onslaught of host immunity (*40*). Indeed, the most advanced efforts at inducing immunity against pre-erythrocytic parasite stages have targeted the sporozoite in the skin (*22*). Despite this, the skin stage of the sporozoite journey remains poorly understood and the effects of the dermal microenvironment and cell traversal on the transcriptional activity of the sporozoite have yet to be explored. To capture this elusive transcriptome, we allowed salivary gland sporozoites to migrate through Transwell μ-pores coated with a monolayer of primary skin fibroblasts, before collecting sporozoites that had either traversed or simply contacted the cell monolayer (Fig. 4A). Whilst our clustering and differential gene expression analysis revealed that sporozoites adapted their transcriptional activity to cues from the host-like microenvironment, medium-only controls were sufficient to induce a gradual shift in gene expression and neither contact with dermal fibroblasts nor cell traversal was found to have an additional impact (Fig. 4B, C). The observed changes (Table S1) were not as wide-ranging as the documented shifts in gene expression experienced by sporozoites exposed to host-like conditions for prolonged periods of time (*41*) or to human hepatocytes (*42*). This suggests that host-induced changes in transcription may occur gradually in a sequential fashion. After one hour, most salivary gland sporozoites had already downregulated the expression of several genes known to be involved in cell traversal (GEST, SPECT1, PLP1, TLP) (*39, 41, 43, 44*). Concomitantly, a small cohort of genes previously implicated in protein folding (*45*) and export (*46, 47*), including HSP70/90, PTEX150 and PV1, was upregulated (Fig. 4D, E, Table S1). PTEX150 is a core component of the *Plasmodium* translocon of exported proteins (PTEX) complex (*46*), responsible for protein trafficking into the host cell following merozoite invasion, while PV1 is a translocon accessory protein that interacts with proteins destined for transport across the translocon (*47, 48*). That the expression of these export machines builds up so rapidly upon initial contact with the mammalian microenvironment, within a timeframe when most sporozoites are still migrating in the skin, suggests that protein export to the host cell might be one of the earliest and most critical events following productive invasion of a hepatocyte. This would mirror merozoite invasion, where the PTEX complex is inserted into the newly formed parasitophorous vacuole membrane within minutes (*49*), enabling export of effector proteins into the erythrocyte to build a suitable niche for replication and immune evasion (*50*). It is worth noting that this activated phenotype was also observed in a small proportion of salivary gland sporozoites before their ejection from the mosquito (Fig. 4B, E). Whether this is the result of spurious signalling events that disrupt sporozoite latency, or instead represents a developmentally regulated process that renders sporozoites transmission-ready awaits to be established.

**Figure 4.**
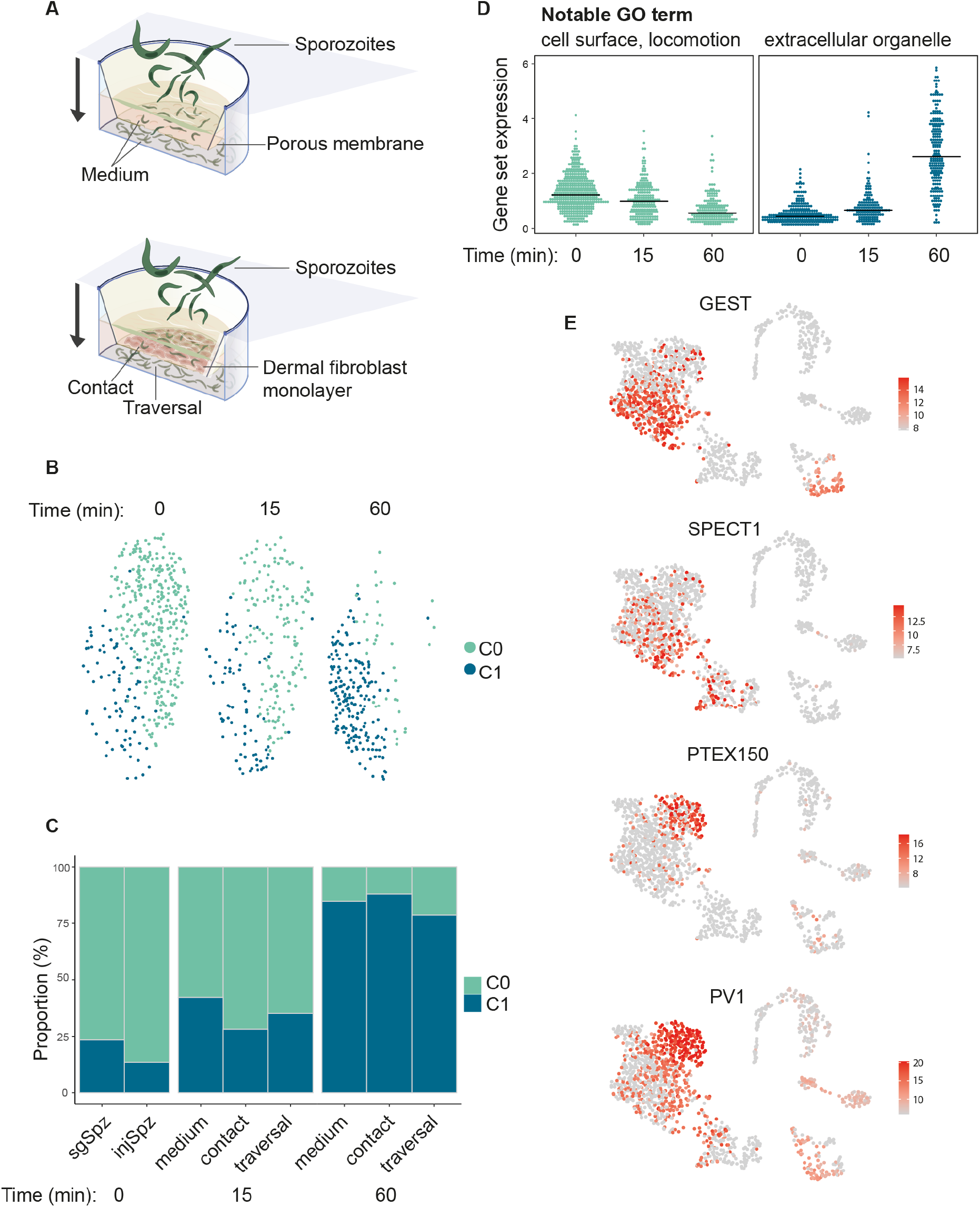
Sporozoite activation in the mammalian microenvironment. **(A)** Schematics of the Transwell sporozoite activation assay. The skin environment was reproduced by transferring sporozoites to Transwell inserts filled with fibroblast growth medium and coated with human primary fibroblasts or kept cell-free at 37°C. Sporozoites were allowed to interact with this setup for 15 or 60 minutes, after which they were collected from both the top and bottom compartments of the Transwell. To reach the bottom compartment in the presence of the fibroblast monolayer sporozoites had to deploy their cell traversal activity, as the Transwell pores were blocked by the tight cell monolayer. ‘Medium’ refers to sporozoites exposed to medium alone, ‘contact’ refers to sporozoites that were collected above the cell monolayer, and ‘traversal’ refers to sporozoites that passed through the cell monolayer and were collected from the bottom compartment. Salivary gland sporozoites kept in insect medium and injected sporozoites were considered as time 0. **(B)** UMAP representations of sporozoite transcriptomes coloured by cluster assignment before and after activation. **(C)** Quantification of sporozoite activation based on the cluster assignments in **B**. **(D)** Average expression of down- and up-regulated genes during sporozoite activation. Each panel comprises all DE genes that present a fold change of at least 1.5 between C0 and C1 and are expressed in more than 25% of the cells (Table S1). The top GO terms in each module are indicated (GO:0009986, GO:0040011, GO:0043230; Bonferroni adjusted *p*-value < 0.01). **(E)** UMAP plots of the complete life cycle with the expression of representative genes from **C** shown. Expression levels are colour-coded as per the scale next to each plot. GEST and SPECT1 are cell traversal genes previously associated with sporozoite quiescence in *P. vivax (41*). PTEX150 and PV1 facilitate protein export to the host cell in blood stage parasites (*46, 47*).

Here we present a comprehensive single-cell analysis of the transcriptional journey of the human malaria parasite *P. falciparum* through its transmission cycle. The transcriptomes presented here provide an in-depth characterization of gene expression across distinct phenotypic stages of development and add a new chapter to the Malaria Cell Atlas, which is an interactive data resource supporting the exploration of individual *Plasmodium* parasite transcriptomes now across the life cycle from two malaria parasite species. This new data set captures key aspects of parasite development within the mosquito and beyond, providing novel insights into how those developmental processes may be regulated, as well as elucidating new potential targets for disrupting the transmission cycle to control malaria. Exploration of correlated gene expression trajectories within this data set can be the starting point to discover the function of genes that lack functional annotation or reveal new and unexpected potential functions for other more well-studied genes. For example, our finding of a possible role for MSP1, a vaccine candidate implicated in merozoite egress, in the release of sporozoites from the oocyst into the mosquito hemocoel, suggests that a conserved egress machinery may be deployed at multiple points in the life cycle. The global view of gene usage across the life cycle made possible by the Malaria Cell Atlas sheds light on core functional strategies that will help inform the development of previously unappreciated multistage antiplasmodial interventions. Such multistage functional classes represent attractive targets for drug development or other interventions aimed at both the host and the mosquito vector.

Transmission of parasites between humans and mosquitoes represent the major population bottlenecks in the malaria life cycle (*1*). After an infectious blood meal, developmental transitions within the mosquito midgut are accompanied by a steep reduction in parasite numbers (down to the tens or units). At the other end of the transmission cycle, not only are the number of sporozoites injected into the skin during a bite generally low, but an even smaller number find their way out of the site of injection and reach the liver to initiate a new phase of population expansion within a liver-resident hepatocyte (*40*). The factors that determine whether an individual parasite will survive this journey are unclear, but one intriguing possibility is that the ability to interact with and respond to the host environment in a productive manner could vary from cell to cell. Of key importance, we identify developmentally regulated transcripts that show highly variable expression across cells of the same stage and could theoretically account for differential phenotypes within a population. Phenotypic validation may reveal whether there is a transcriptional signature that correlates with infectiousness and predicts the likelihood of successful transmission. Further interrogation of this data set will serve as a major accelerator for the discovery of new targets to block human malaria transmission and ultimately prevent establishment of malaria disease.

## Materials and Methods

### Parasite maintenance and isolation

#### P. falciparum maintenance and gametocyte induction

*P. falciparum* gametocytes were induced and cultured as before (*51*). Briefly, asexual blood stage cultures were maintained in asexual culture medium (RPMI 1640 with 25 mM HEPES (Life Technologies), 50 μg L-1 hypoxanthine (Sigma), 0.3 g L-1 L-glutamine (Sigma), and 10% A+ human serum (Interstate Blood-Bank)) with an hematocrit (HC) of 5%. Gametocyte cultures were induced at 2.5% parasitemia and maintained for 16-17 days in gametocyte culture medium (RPMI 1640 with 25 mM HEPES (Life Technologies), 50 μg L-1 hypoxanthine (Sigma), 2 g L-1 sodium bicarbonate (Sigma), 0.3 g L-1 L-glutamine, 5% A+ human serum (Interstate Blood-Bank) and 0.5% AlbuMAX II (Life Technologies), 5% HC) with daily media changes.

#### Standard membrane feeding assay

*Anopheles stephensi* mosquitoes were reared under standard conditions (26°C-28°C, 65%-80% relative humidity, 12 hour:12 hour light/darkness photoperiod). Adults were maintained on 10% fructose, except for the 12 hours before an infectious blood meal, when mosquitoes were starved. Mosquito infections were carried out using a standard membrane feeding assay (SMFA), as previously described (*52*). In brief, 15-17 day-old gametocyte cultures with at least 2% stage V gametocytaemia were diluted in a 50% mixture of fresh RBCs and A+ human serum and then used to feed starved female mosquitoes. Mosquitoes were allowed to feed for 12 minutes, after which they were kept without fructose for 24 hours, so that only blood-fed mosquitoes would survive. From then on, mosquitoes were given fresh fructose daily until the day of dissection.

#### Retrieval of gametocytes

At day 16 post induction, gametocytes were loaded on a 70% percoll gradient and spun at 500g for 10 minutes without brake at 38°C. Gametocytes were washed once in gametocyte medium at 38°C and the pellet resuspended in 1 ml of suspended animation (SA) buffer (*53*) supplemented with 2 μg/ml Hoechst 33342 and 4 μg/ml Pyronin Y. In the presence of Hoechst 33342, interactions between Pyronin Y and DNA are disrupted and Pyronin Y mainly stains RNA (*54*). Gametocytes were stained at 4°C for 20 minutes. After 20 minutes, cells were washed twice in SA and resuspended in fresh SA, passed through a strainer with single gametocytes sorted into a 96-well plate using a BD FACsAriaIII at 4°C. PyroninY was detected using a 561nm laser with a 586/16nm filter, and Hoechst 33342 was detected using the 488 nm laser with the 450/50nm filter (Fig. S2). Gametocytes were collected from parasites deriving from either an NF54 genetic background (African) or a recent, culture-adapted field isolate, APL5G (Cambodian)(*55*).

#### Retrieval of ookinetes

24 hours after SMFA, mosquitoes were dissected, and the blood bolus removed from the midguts. The collected blood boli were placed into ice-cold PBS supplemented with 2x protease inhibitors (Halt™ Protease Inhibitor Cocktail, Thermo Fisher) and 50mM EDTA. For the first pilot flow cytometry experiment (Fig. S1A), uninfected RBCs and activated gametocytes cultures were placed in ookinete medium (RPMI-HEPES supplemented with 100uM xanthurenic acid, 200 uM hypoxanthine and 10% BSA, pH7.4) for antibody- and DNA staining, and used as controls. Each sample was split into four aliquots, one aliquot remained unstained and the others were incubated with either 1:500 anti-Pfs25-cy3 antibody (*51*), 1:1000 DRAQ5 DNA stain (BioLegend), or both, respectively, at 4°C in the dark for 30 minutes. Samples were then washed twice in ice-cold PBS and resuspended in ice-cold PBS, before being analysed on a BD Fortessa using the 561nm laser and filter 586/15 for cy3 and the 640nm laser and 730/45 filter for DRAQ5, respectively. For the sorting, ookinetes collected 24 and 48 hours after a feed were prepared as above and sorted into a 384-well plate. One-fifth of the plate was sorted from the P1 population and rest from the P2 population (Fig. S1B, S1C). Oocysts per midgut were counted on day 10 post-feed from 41 mosquitoes and revealed a prevalence of infection of 63%.

#### Retrieval of sporozoites

Sporozoites were collected from different anatomic sites in the mosquito to capture different stages of development. Before dissections, mosquitoes were cold anesthetized (4°C) and killed inside a secure glove box by beheading. Oocyst and salivary gland sporozoites were obtained by dissecting mosquito midguts and salivary glands, 12 and 17-20 days post blood meal, respectively. Hemolymph sporozoites were released from mosquito carcasses on day 14 after removal of both salivary glands and midguts. All samples were initially homogenised with a disposable pestle to release sporozoites from the mosquito tissues. After a filtration step with a 20 μm filter (Pluriselect), sporozoites were loaded onto a 3 ml Accudenz cushion and centrifuged at 2,500 g for 20 min (*56*). All the above procedures were carried out at 4°C in Schneider’s insect medium. After centrifugation, sporozoites were collected from a thin layer on top of the Accudenz cushion (mosquito debris separated to the bottom), washed once with cold Schneider’s buffer (12,000 g, 3 min, 4°C) and resuspended in the same buffer supplemented with 1% FBS. Before FACS sorting, sporozoites were immunostained for 30 minutes with anti-CSP (2A10 clone, MR4) diluted at 1:200, followed by a 20 minute incubation with an Alexa Fluor 488-conjugated antimouse antibody (1:300, Jackson ImmunoResearch) and DRAQ5 (1:1000, BioLegend). All incubations and washes were carried out at 4°C. Single cells were sorted directly into NEB lysis buffer on a BD FACSAria III equipped with a 100 um nozzle.

Sporozoites released through a mosquito bite were collected by allowing mosquitoes to feed on a warm fructose solution (80 g/L) supplemented with 10% serum for 5 minutes. After confirming that sporozoites had been released and that no mosquito debris were present in the sample, sporozoites were stained for 20 min with Draq5 at 4°C and sorted directly into NEB lysis buffer.

#### Cell traversal assay

Sporozoites released from salivary glands on day 20 after a blood meal were purified as before and resuspended in fibroblast growth medium (PromoCell) supplemented with heat-inactivated bovine serum (Sigma) to a final concentration of 10%. 50,000 sporozoites were added per well of a 24-well plate fitted with Transwell inserts (5 um pore size, Corning) coated with 100,000 human primary dermal fibroblasts (PromoCell) to form a tight monolayer. Sporozoites were allowed to migrate for 15 or 60 minutes and were then recovered separately from the top and bottom Transwell compartments. On average, 35% of the input was recovered from the bottom Transwell compartment. Some control wells did not contain fibroblasts, so that the effect of the medium on the sporozoites could be evaluated. Before FACS sorting, sporozoites were immunostained as described above, with all steps carried out in fibroblast growth medium with 10% FBS at 4°C. Single cells were sorted as before.

### scRNA-seq

Cells were sorted in 96-or 384-well plates on FACSAria III into NEB lysis buffer (0.1x cell lysis buffer E6428B-SG, 0.05x RNAse-inhibitor Murine E6429B-SG and 0.85x Nuclease-free water E6433B-SG, from New England Biolab). The ookinetes were sorted into a 384-well plate, and all other samples were sorted into 96-well plates. The 384-well plate contained 2 μL lysis buffer, whereas the 96-well plates contained 4 μL. Reverse transcription and PCR was performed in the Sanger Institute single-cell sequencing pipeline using the NEBNext Single Cell/Low Input RNA Library Prep Kit for Illumina (E6420L) with 26 PCR cycles. cDNA was purified using an AMPure XP bead clean-up on either an Agilent Bravo or Hamilton Star. Reaction volumes were doubled in line with the starting volume of the lysis buffer in 96-well plates for cDNA generation, amplification and purification. The samples were quantified using the Mosquito and Omega plate reader to calculate volume of cDNA needed for library preparation. Libraries were made using the scRNA cDNA-XP plates, bravo platforms and NEB Ultra II FS reagents (E7805L). Libraries had an additional eight cycles of PCR. Libraries were then pooled to 384 samples prior to AMPure XP bead Clean-up on a Hamilton Star. The samples were run on an Agilent Bioanalyser and normalised to 32.8 nM prior to sequencing. Samples were sequenced on a HiSeq 2500 with 75 bp paired-end reads.

### scRNA-seq mapping and analysis

#### Single-cell transcriptome mapping and read counting

Data acquisition and mapping followed the pipeline described in (*6*). Briefly, reads were trimmed using trimgalore (cutadapt v 1.18) (*57*). Trimmed reads were mapped to the *P. falciparum* v3 genome (https://www.sanger.ac.uk/resources/downloads/protozoa/, October 2016) with HISAT2 (v 2.1.0) (*58*) using *hisat2 --max-intronlen 5000 p 12*. Mapped reads were then summed against transcripts with featureCounts in the Subread package (v 2.0.0) (*59*) using *featureCounts -p -t CDS -g transcript_id*.

#### Quality control and normalisation

Given the high variability of genes detected across the life cycle (*8*), low quality cells were identified based on the distribution of the number of reads and genes detected per cell within each parasite stage (gametocytes, ookinetes, sporozoites). For gametocytes, cells with fewer than 500 genes and 10000 reads per cell were removed. For ookinetes, cells with fewer than 400 genes and 5000 reads per cell were removed. For sporozoites, cells with fewer than 40 genes per cell and 5000 reads per cell were removed (Fig S4, Table S1). Transcriptomes were normalised using a deconvolution size factor with scran (v 1.16.0) (*60*) using quickCluster() and calculateSumFactors() to account for the large differences in detection between parasite stages. Visual inspection of the cell-wise relative log expression plots (Fig S4) shows normalisation by scran smoothed profiles across stages.

#### Cell clustering and projection

To delineate parasite transcriptomes into cell-types we performed graph-based Louvain clustering in Seurat (v 3.1.5) (*12*) with different resolution depending on the analysis (overview of all cells and integration: resolution = 0.5; sporozoite and ookinete development: resolution = 0.6; sporozoite activation: resolution = 0.4). Transcriptomes were visualized using UMAP nonlinear dimensionality reduction in either Scater (v1.16.0) (*61*) with n_neighbors=5, min_dist=1, spread=3 (global analysis, Figure 1) or Seurat with n.neighbors=5, min.dist=2, and spread=3 (integration, Figure 2) or default parameters (all other analyses).

#### Gene graph and clustering

To create the k-nearest neighbor (kNN) gene graph, cells from cluster 0 and 1 (sporozoite clusters) were downsampled to 100 cells each in order to get a more even representation across cell-types. Feature selection was performed on the expression matrix of these 623 transcripts to identify the top 2000 biologically-relevant genes using selectFeature() in scmap (v 1.10.0) (*14*). Transcripts were further filtered by removing those with fewer than 200 total reads across the 623 cells resulting in 1797 genes that were used to generate the gene graph. The counts matrix was normalized by gene by dividing the mean counts for each gene and log scaling in order to reduce the amount in which clusters were driven by total expression level. The kNN graph was made using this gene normalized expression matrix with the Nearest Neighbors subpackage in python’s scikitlearn (v 0.2.22.post1) with n_neighbors=6 and a manhattan distance metric (*62*). Spectral clustering was then performed on the kNN graph using the SpectralClustering subpackage of scikitlearn (v 0.2.22.post1) (*63, 64*). The kNN graph was then rendered in Gephi with force atlas 2 layout algorithm in linlog mode with scale of 0.2 and gravity of 0.8 (*65*). To test for functional class enrichment in each gene cluster, a Gene Ontology (GO) analysis was conducted using the GO analysis tool on PlasmoDB. Pearson’s correlations between ApiAP2 TFs expressed across the data set and each gene cluster were computed using the cor function in R.

#### Integration with P. berghei data

To identify conserved and divergent patterns of expression in *Plasmodium* transmission stages, we compared our *P. falciparum* data to *P. berghei* data from (*8*). To do this, we first subsetted similar stages across the two datasets. (*P. falciparum:* gametocytes, ookinetes, and all sporozoite collections; *P. berghei:* gametocytes, ookinetes, and gland and injected sporozoites). We then built expression matrices for each dataset based on one-to-one orthologs as described in (*8*). Using these orthologous expression matrices, we integrated the two datasets using Seurat’s integration function by first identifying anchors with the FindIntegrationAnchors() and then using IntegrateData() (*12*). Cells were visualized and clustered based on the integrated expression values as described above. Conserved markers and differential expression between species was then performed on each cluster using the FindConservedMarkers() and FindMarkers() functions in Seurat (*12*).

#### Pseudotime

Ookinetes and sporozoites were ordered along two independent developmental trajectories using Slingshot (*66*). In brief, Seurat clusters and UMAP cell embeddings were used as input to build a minimum spanning tree to infer the order of the clusters. Slingshot then fits principal curves to estimate pseudotime values for each cell. In both cases, Slingshot identified a single trajectory that described the progression of the cells along developmental pseudotime. To identify genes differentially expressed as a function of pseudotime, we used the differentialGeneTest function in Monocle 2 (*67*). DE genes were clustered hierarchically using the pheatmap package with a Pearson correlation distance measure (clustering_distance_rows = “correlation”). Enrichment of GO terms in each pseudotime cluster was tested using the GO analysis tool on PlasmoDB. To assess whether developmentally-regulated genes showed variable expression between individual cells of the same developmental stage, we identified highly variable genes (HVG) using M3Drop (*68*). For sporozoites, ooSpz, hlSpz, sgSpz, and injSpz cells were analysed separately. For the analysis of HVGs in ookinete cells, we first used a general linear model to regress out the effect of pseudotime. Genes with high variability across cells were then identified using M3Drop, as previously described (*8*). HVGs with a q-value < 0.05 were then intersected with the pseudotime-regulated gene subset. Pearson’s correlations in Fig. 3H and Fig. S6F were computed using the cor function in R.

#### Motif discovery

To find short DNA motifs enriched in the gene clusters from **Fig. 1D**, the 1000 bp upstream of the start codon of each gene in each cluster were analysed with DREME (*69*). The negative data set included the equivalent 1000 bp region of all genes outside the cluster being analysed. Motifs were considered enriched if the e-value was lower than 0.05. Tomtom (*70*) was used to compare the top scoring motif of each cluster with transcription *cis*-regulatory elements described in *P. falciparum* (*71*–*74*).

#### Comparison with sporozoite bulk transcriptomes

Sporozoite single cells were assigned one of two stages (ooSpz or sgSpz) based on gene expression correlations between scrNAseq (this study) and bulk (*25*) transcriptomes. Pearson and Spearman correlations were computed using the corr package in R. To determine whether a transcript was likely to be translationally repressed (TR), we intersected scRNAseq gene expression profiles along pseudotemporal development with the published proteomes of ooSpz and sgSpz (*25*). Sporozoite transcripts that were expressed in our data set but absent in the proteome of the same stage were labelled as TR (Table S1).

## Supporting information

GeneTable

CellInfo

Counts

EnrichedMotifs

GOTerms

## Supplemental Figures

**Figure S1.**
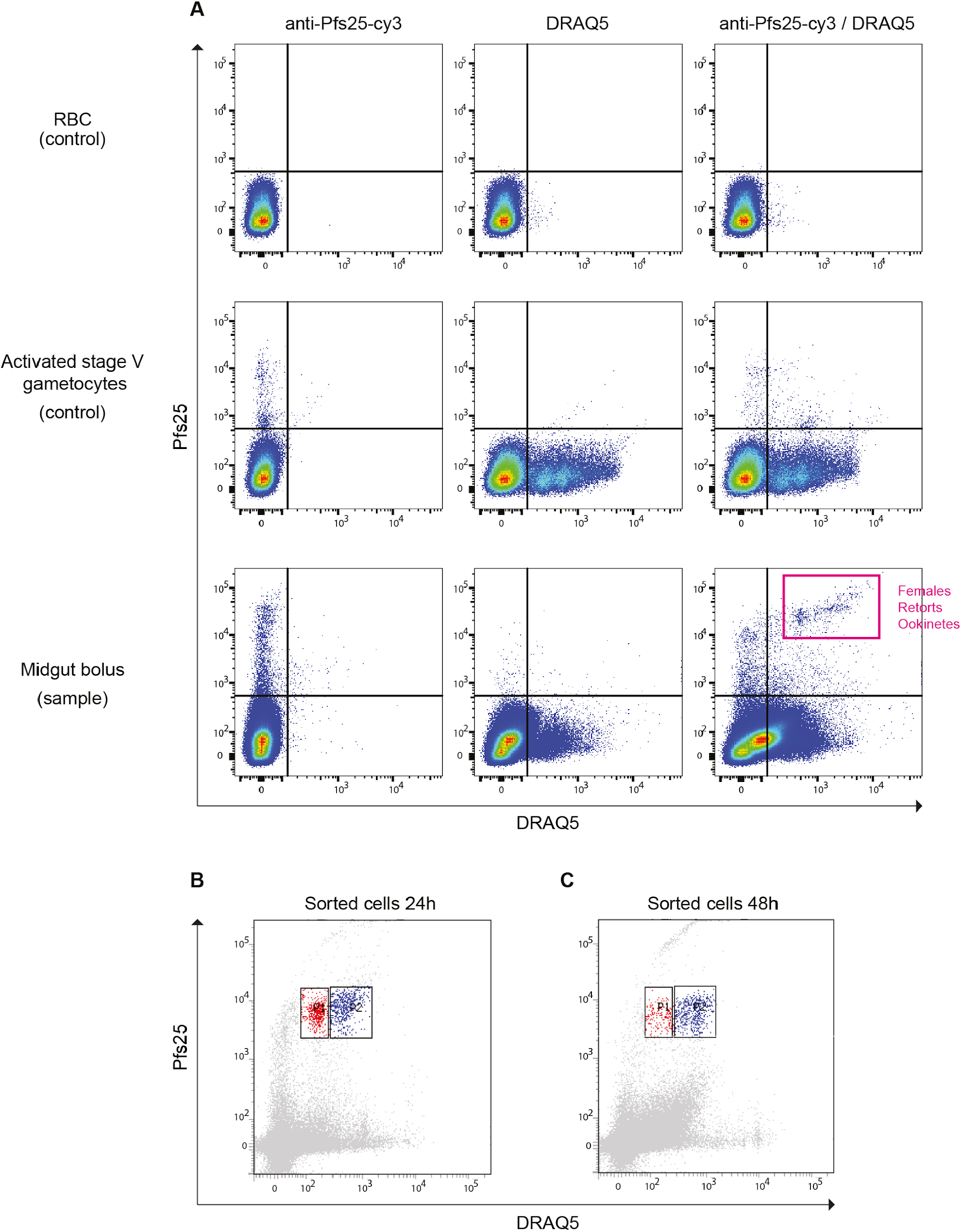
Retrieval of *P. falciparum* ookinetes. **(A)** Pilot flow cytometry experiment using the DNA-marker DRAQ5 and the anti-Pfs25 antibody coupled to cy3, to identify *P. falciparum* ookinetes. Uninfected RBCs and activated stage V gametocytes were used as controls. Double-positive cells, which comprise activated female gametes, retorts and ookinetes, were collected from *Anopheles stephensi* midgut boli 24 hours after an infectious blood feed. **(B-C)** FACS scatter plots showing the two populations (P1 and P2) that were collected for the 24 h **(B)** and 48 h **(C)** post-feed timepoints.

**Figure S2.**
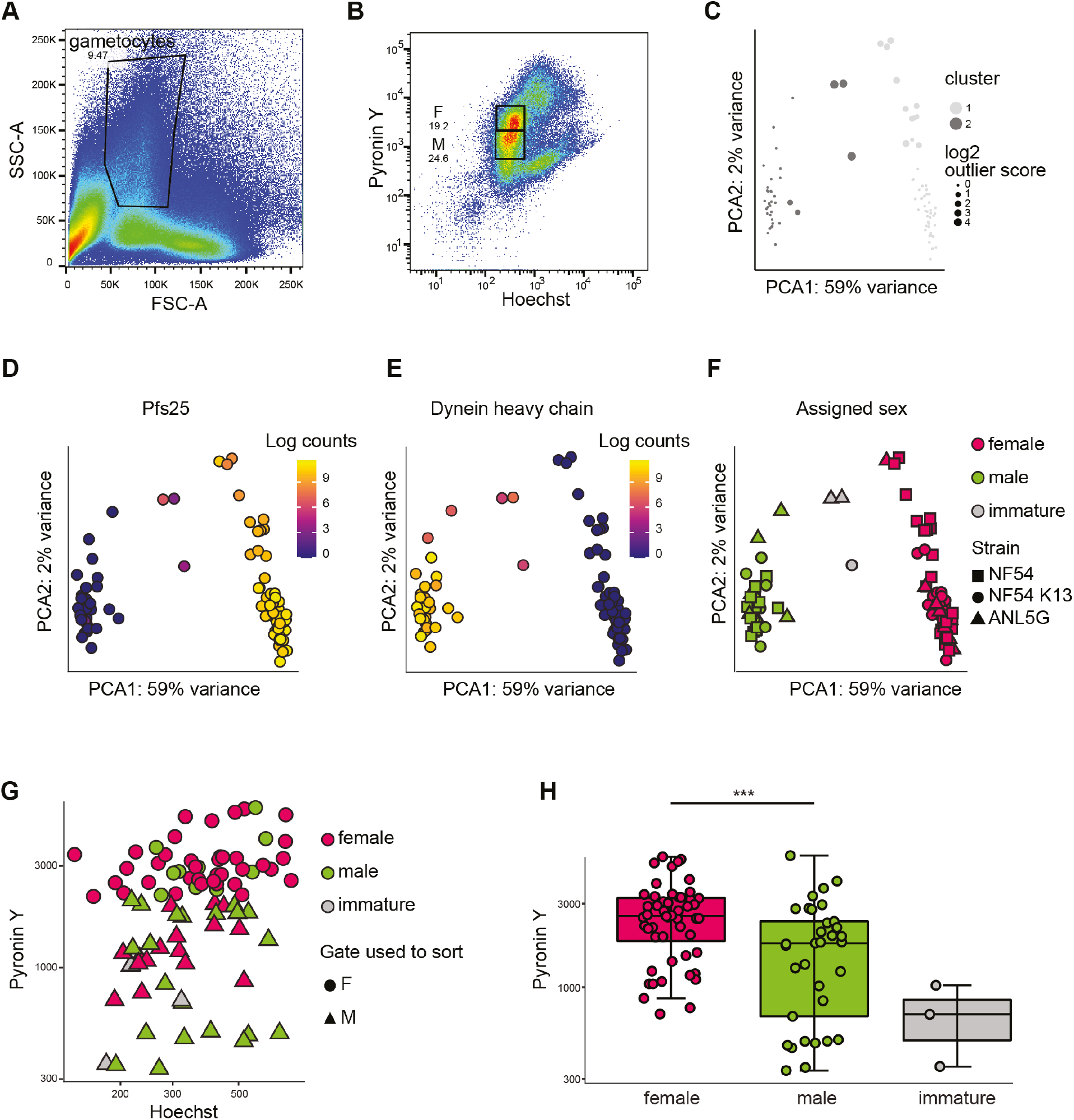
Use of DNA/RNA double staining to sort *P. falciparum* gametocytes by sex. **(A)** Scatter plot showing Forward (FSC) and Side-scatter (SSC) of an unpurified 15 day-old gametocyte culture. **(B)** Scatter plot showing the population gated in **A** stained with DNA (Hoechst) and RNA (Pyronin Y) dyes. Gates around the two cell populations used for single cell sorting are shown. Gates are labelled F for females and M for males. (**C)** Cluster analysis of the sorted gametocyte populations. Each dot represents one gametocyte. Dot sizes represent the outlier score of the cluster assignments. **(D-E)** Principal component analysis (PCA) of gametocyte single cell transcriptomes, highlighting expression of Pfs25, a female-specific marker **(D)** and dynein heavy chain protein (PF3D7_0905300), a male-specific marker **(E)**. **(F)** PCA showing the assigned sex for each cell, based on data from C-E. The different shapes represent the three different parasite lines sorted, two deriving from the canonical lab strain NF54 background, with a third deriving from a recently culture-adapted Cambodian field isolate, APL5G. Gametocytes from the three parasite lines group together by sex on the PCA, reflecting the similar transcriptional profiles between strains. **(G**) RNA and DNA content in sorted gametocytes. Cells are coloured by their assigned sex and shaped according to the gate used for sorting. RNA and DNA stains are indicated in the y- and x-axis, respectively. **(H)** Box plots summarising the findings from G. Female gametocytes are significantly enriched by pyronin Y stain compared to male gametocytes (Welch’s t-test, p-value = 4.873e-05), indicated by asterisks.

**Figure S3.**
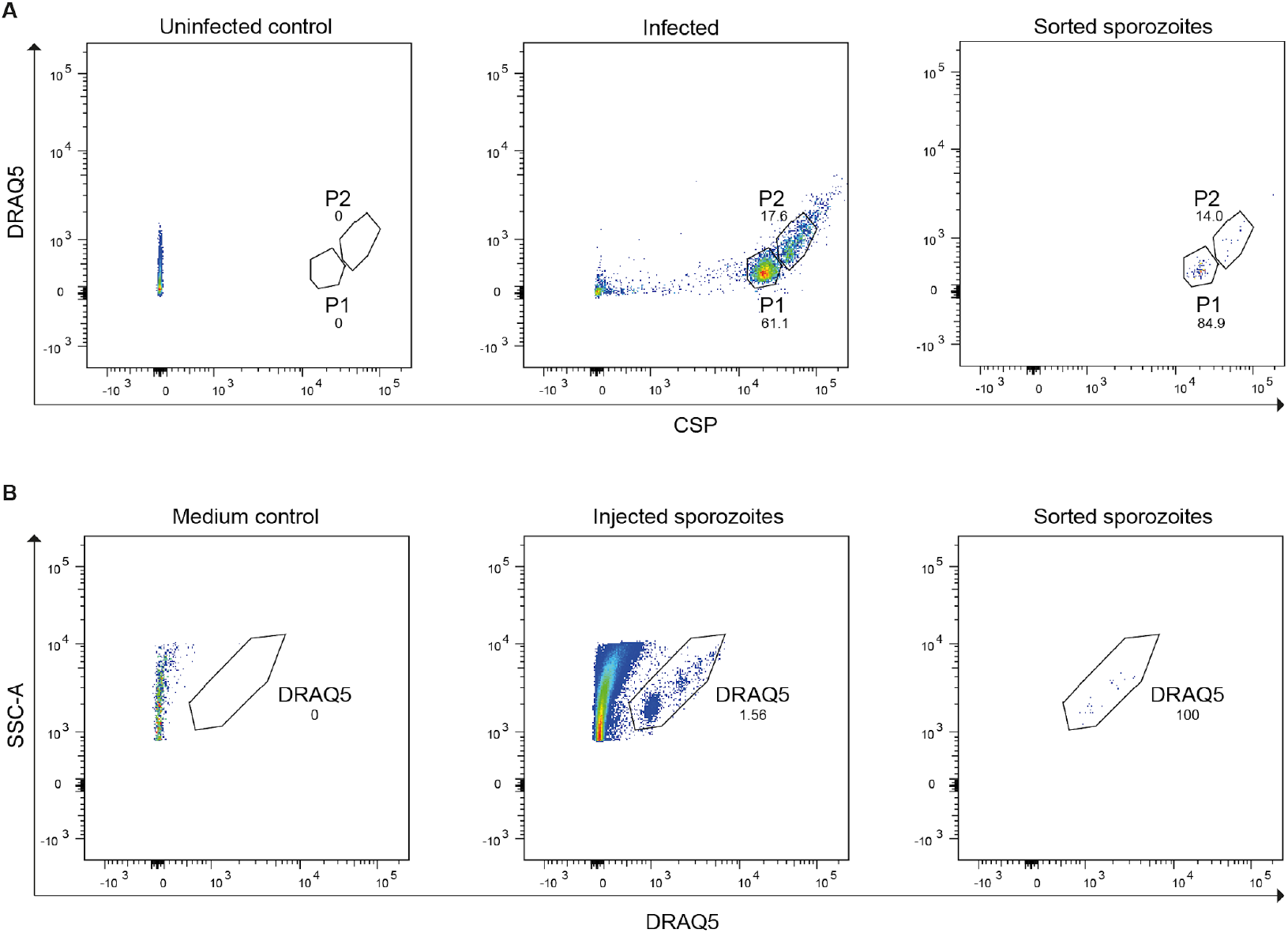
Retrieval of sporozoites. **(A)** Gating strategy for sorting sporozoite cells. Partially purified cells (see Materials and methods) were stained with the DNA-stain DRAQ5 and anti-CSP antibody to sort sporozoites from mosquito debris. Most cells were collected from gate P1, where 80% of all sporozoites formed a compact population. We also collected a smaller number of cells from the more scattered P2 population, but we subsequently confirmed by qPCR using CSP-specific primers that P2 contained a much higher proportion of *non-Plasmodium* cells than P1, which was 95% positive for CSP by qPCR (not shown). **(B)** Sporozoites released through a mosquito bite were sorted based on the DRAQ5 signal alone, as no mosquito debris was detected in the sample either visually or by qPCR.

**Figure S4.**
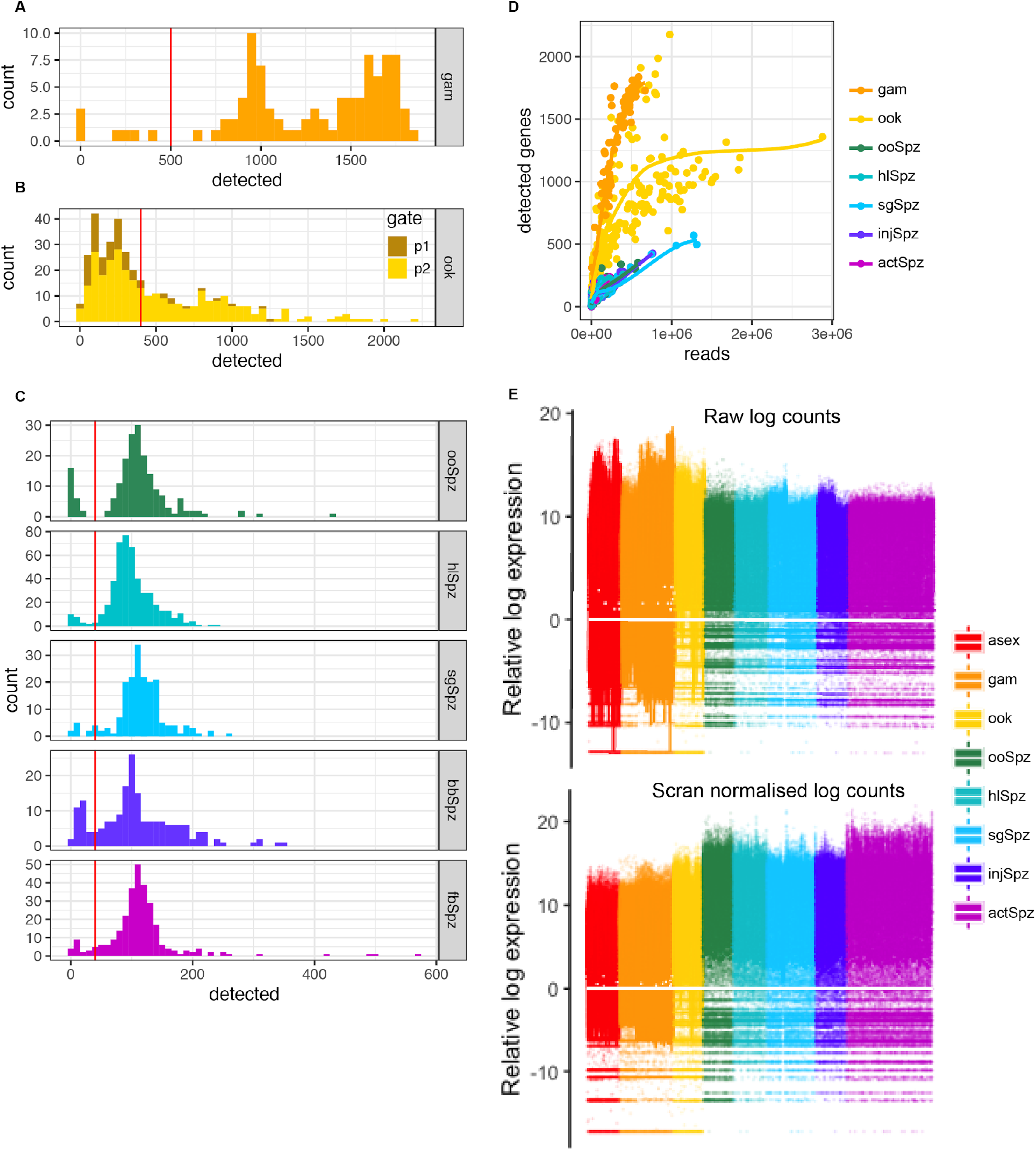
Quality control and normalisation. Filtering of cells was based on the distribution of the number of genes per cell within each parasite stage. **(A)** The distribution of genes detected in gametocytes. Cells with fewer than 500 genes were removed. **(B**) The distribution of genes detected in ookinetes. Cells with fewer than 400 genes were removed. Cells are colored according to the sorting gate (Fig S1). The majority of cells from p1, which had lower DNA content, were excluded based on this threshold, supporting the hypothesis that these may be dead or dying unfertilised macrogametes. **(C)** The distribution of genes detected in sporozoites (faceted by collection). Cells with fewer than 40 genes were removed. **(D)** The number of reads per cell versus the number of genes detected varied depending on parasite stage. Gametocytes and ookinetes with fewer than 10000 reads were removed. Sporozoites with fewer than 5000 reads were removed. **(E)** Cell-wise relative log expression plots with raw log counts (top panel) and scran normalised log counts. Normalisation somewhat smoothed the heterogeneity seen between parasite stages.

**Figure S5.**
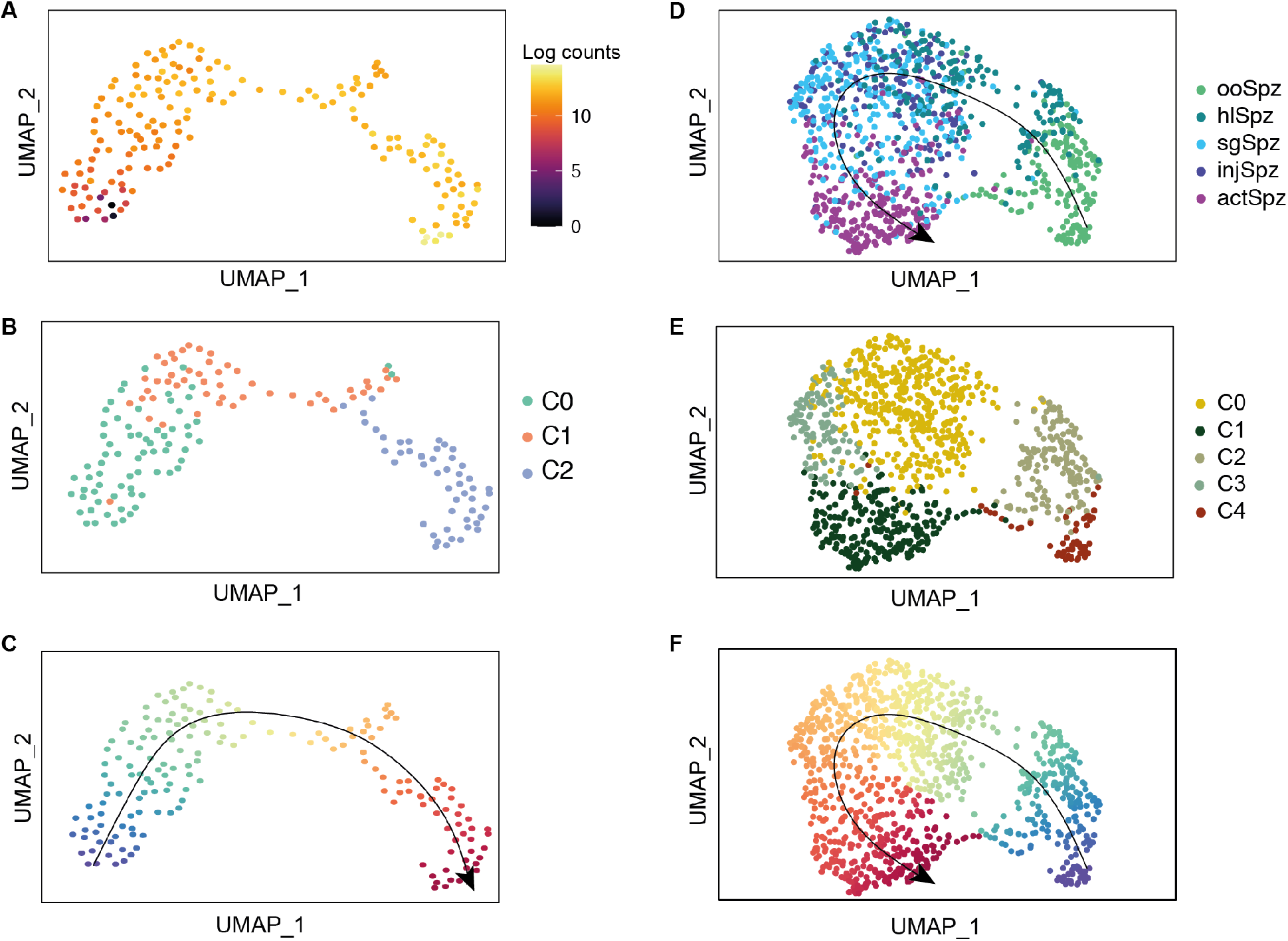
Pseudotime ordering of asexual and sporozoite transcriptomes for gene graph. **(A-C)** UMAP of asexual stage transcriptomes from (*6*) coloured by **(A)** expression of the late-stage marker MSP1 (PF3D7_0930300) (*30*), **(B)** Seurat cluster assignment, and **(C)** pseudotime progression. **(D-E)** UMAP representations of all sporozoite transcriptomes in the data set coloured by sorted stage **(D)**, Seurat cluster assignment **(E)**, and pseudotime **(F)**. The lines in **C** and **F** represent the pseudotime trajectory. Pseudotime values were calculated using the Seurat clusters in **B** and **E** as input to Slingshot and then used to order cells in **Fig. 1E** by their developmental progression.

**Figure S6.**
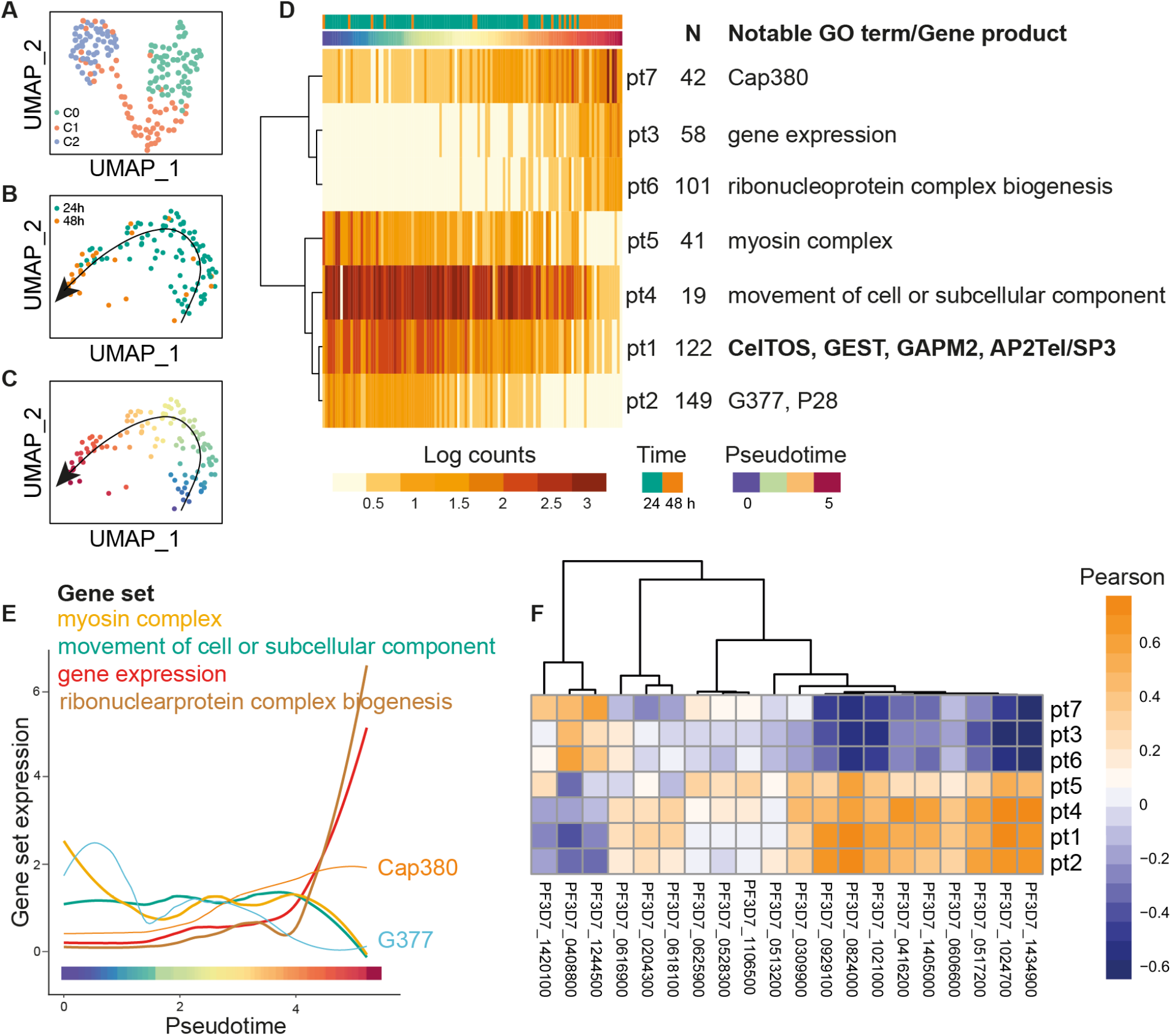
Ookinete development. **(A)** UMAP representation of single-cell transcriptomes from ookinetes coloured by cluster assignment. Cluster 2, which was composed primarily of cells from the 48 hour collection, was excluded from further analysis as it lacked expression of known maker genes. We hypothesize these cells could represent dying ‘ghost’ ookinetes, which are at high prevalence at this time point (*13*). **(B, C)** UMAP plots excluding cells from cluster 2 colored by time post blood meal (**B**) and developmental pseudotime (**C**). We identified 533 DE genes over pseudotime (q-value < 0.001) that clustered into 7 modules (pt1-pt7) with different expression dynamics. **(D)** Heatmap showing the mean expression of each cluster of DE genes over pseudotime with cells ordered by their pseudotime value. The number of genes (N) and top GO terms (Bonferroni adjusted p-value < 0.05) in each cluster are shown. Examples of highly variable genes between individual ookinetes are indicated in bold next to the cluster to which they belong. HVGs were determined using a general linear model to regress out the effect of pseudotime and M3Drop with a false discovery rate < 0.05. Heterogeneity in the expression of genes associated with cell traversal (CelTOS) and motility (GAPM2) suggests phenotypic heterogeneity with respect to midgut colonisation. **(E)** Expression of sets of functionally related genes in each cluster over pseudotime. **(F)** Heatmap of Pearson’s correlations between the expression of 20 species-conserved (*P. berghei* and *P. falciparum*) genes of unknown function and gene clusters from **(D)**. Patterns of correlated expression can help assign roles for genes that lack functional annotation. For instance, expression of PF3D7_1434900, PF3D7_1024700, and PF3D7_0416200 is highly correlated with that of genes involved in cell movement, suggesting a possible role in ookinete motility.

**Figure S7.**
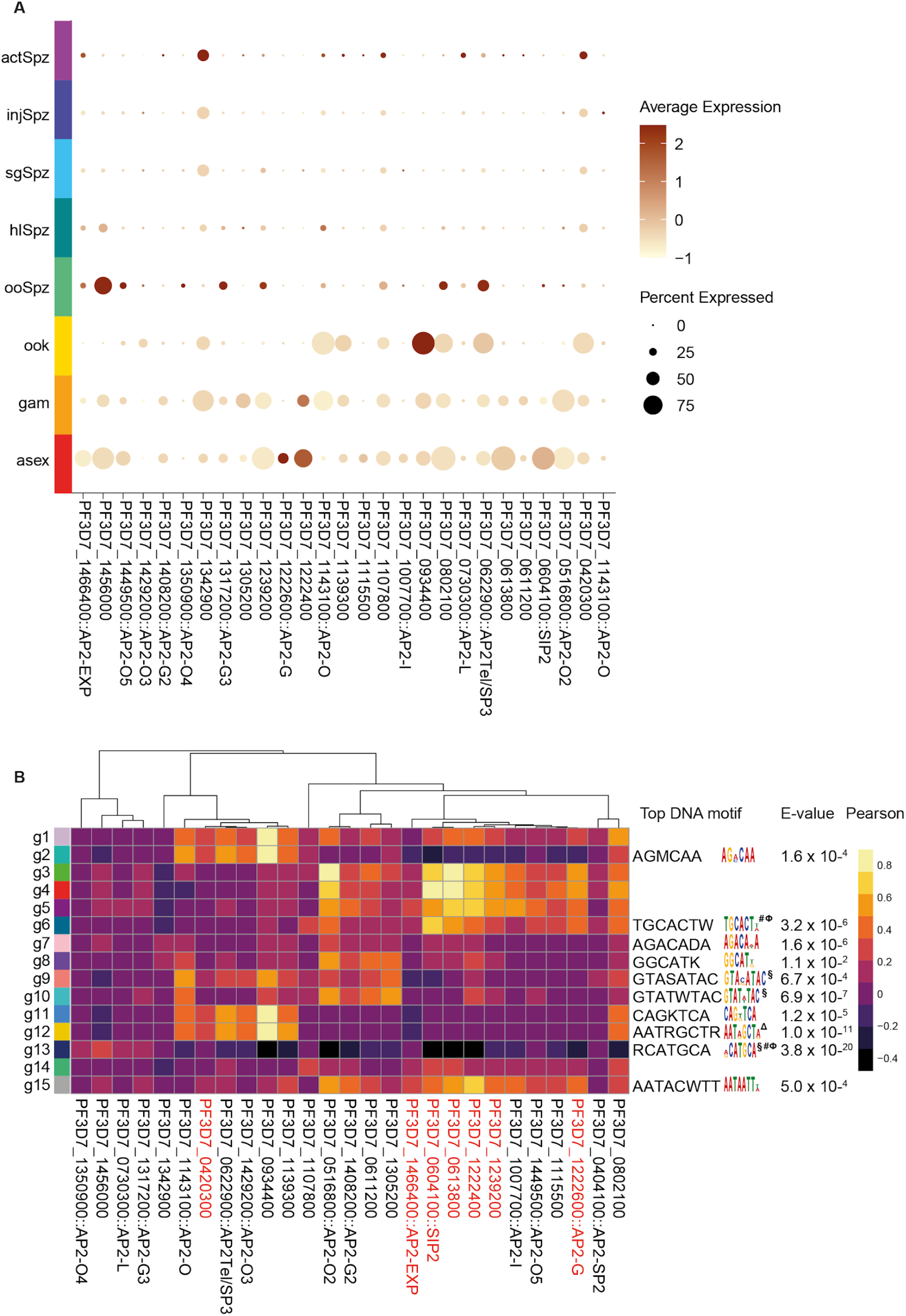
Expression of ApiAP2 transcription factors across the life cycle of *P. falciparum*. **(A**) Dot plot showing the average expression (colour-coded) and proportion of cells of each stage (dot size) that express each of the 26 ApiAP2 TFs detected in the data set. **(B)** Pearson’s correlations (colour-coded) between ApiAP2 gene expression and the gene clusters from figure 1B. The seven ApiAp2 TFs highlighted in red are ApiAP2s for which BDP1 association with their promoter regions has been demonstrated (*18*), suggesting they are subject to epigenetic regulation. Most are strongly correlated with asexual and sexual stages of the life cycle. DNA motifs enriched within 1 Kb upstream of the start codon of genes in each cluster are shown (Evalue < 0.05), except when no statistically significant enrichment was found. Motifs identified in previous studies are marked with § (*72*), # (*73*), Φ (*71*), and Δ (*74*) and show the expected correspondence with the life cycle stage they were linked with. The top motif in cluster g12 corresponds to the AP2-O binding site identified in Δ (*74*), while the most significant motifs in clusters g6 and g13 match the binding sites for SIP2 and AP2-EXP, respectively (*71*). The former has been shown to bind to subtelomeric chromatin in *P. falciparum* and has been implicated in the silencing of upsB *var* genes (*75*), which are represented in cluster g6. Our analysis shows that expression of SIP2 is highly correlated with that cluster, which is consistent with SIP2’s putative role in *var* gene regulation.

**Figure S8.**
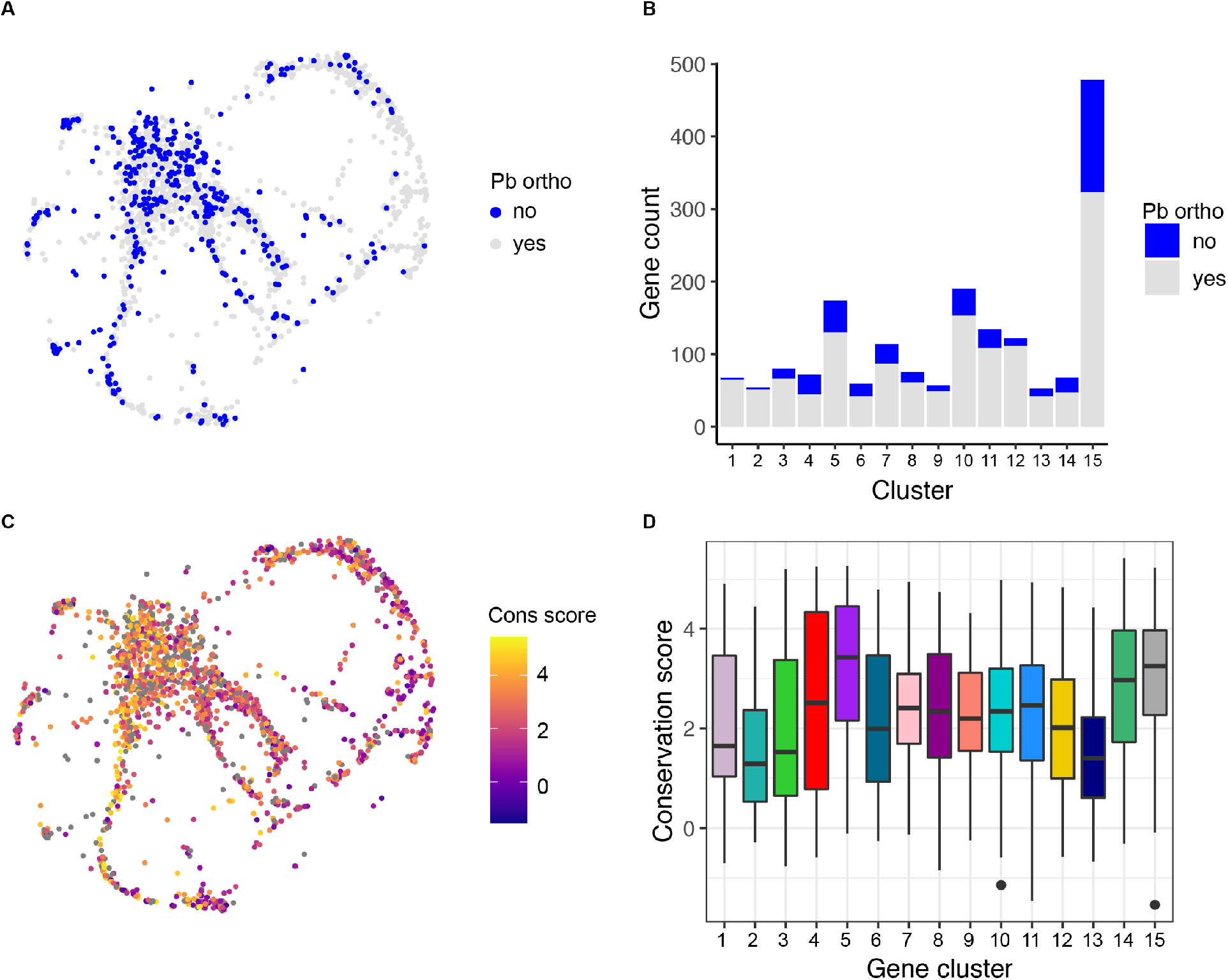
Gene and amino acid conservation scores across the kNN gene graph. **(A)** The kNN gene graph with non-orthologous genes highlighted in blue. **(B)** A barplot of gene counts for each gene cluster (g1-g15) colored by orthology. Cluster g15 was significantly enriched for genes that have no orthologs with *P. berghei*, and clusters g1, g2, g12 had more genes with one-to-one orthologs than expected by chance (Fisher’s Exact Test, bonferroni, *p* < 0.05). **(C)** Amino acid conservation score between *P. falciparum* and *P. berghei* based on mean substitution score for all amino acids in the protein (*6, 76*). (**D**) A boxplot of conservation score by gene cluster. There was significant variation in conservation score across the clusters (ANOVA, *p* < 0.01) with clusters g2 and g13 having the lowest median conservation score. These clusters are also enriched for genes involved in host-parasite interactions.

**Figure S9.**
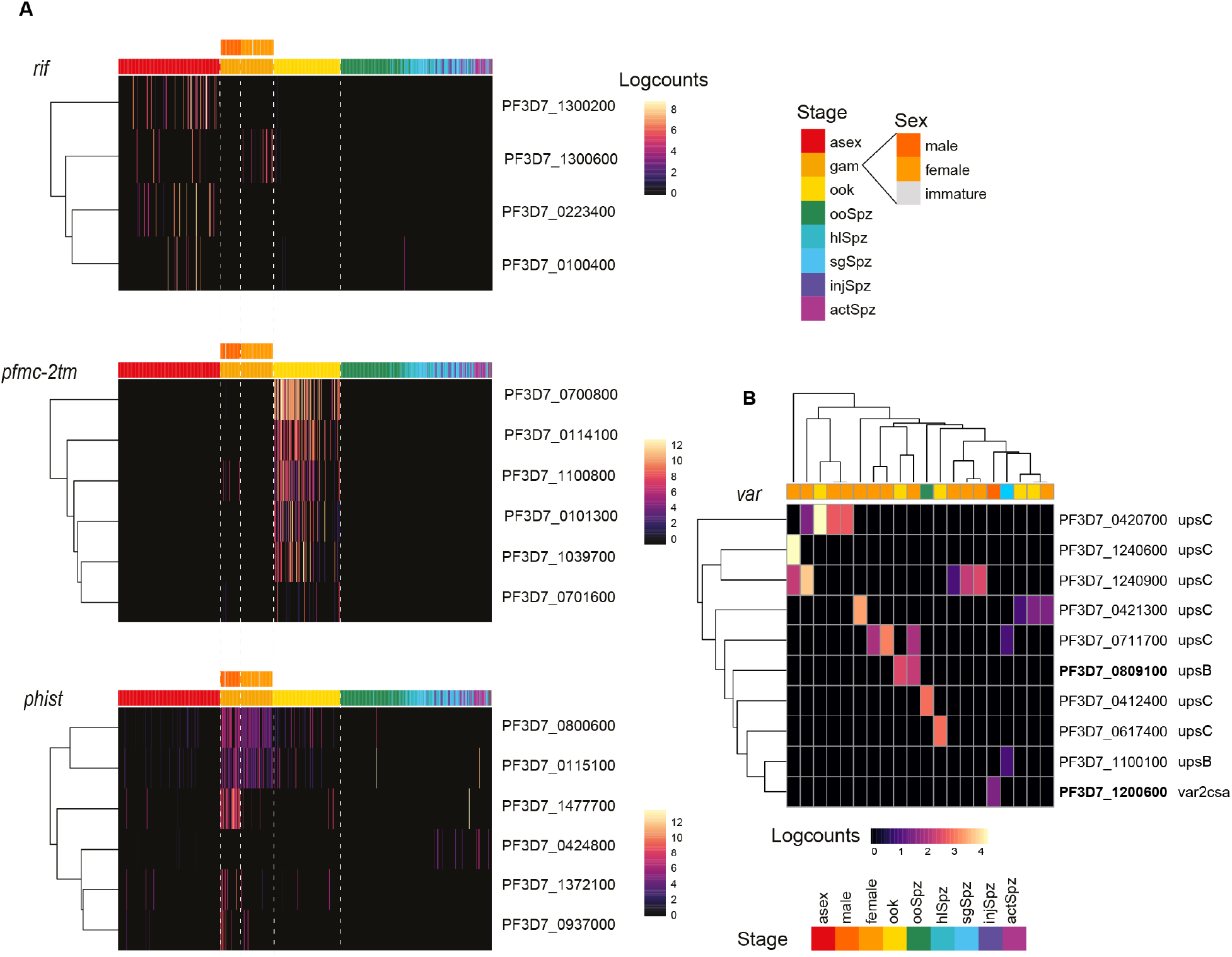
Expression of *P. falciparum* multigene families across the life cycle. **(A)** Heatmaps showing the expression levels of genes without 1:1 *P. berghei* orthologs across the life cycle. Only genes that belong to clonally variant multigene families (*77*) and expressed in at least 10 cells across the data set are shown. Ookinetes are significantly enriched for expression of six genes belonging to the PfMC-2TM family (Bonferroni adjusted p-value = 1.2 x10^−05^), which have been localised to the host-parasite interface in asexual parasite forms (*78*) and were not known until now to be expressed in this stage of the life cycle. Expression of PfMC-2TM family members is associated with ookinete development over pseudotime (q-value < 0.01) (Table S1). **(B)** Expression levels of members of the *var* multigene family for which sense transcripts could be detected. We identified sense transcripts using split reads spanning the var intron as described in (*6*) and focused on these because ncRNA is expressed from the body of members of this gene family. Each column in the heatmap represents a single cell colour-coded by its life cycle stage. The type of each *var* gene type is indicated as upstream promoter sequence (ups) B/C or *var2csa*. Although we were only able to confirm sense transcription of *vars* in a small number of cells, several of the identified transcripts had been previously identified in transmission stages. PF3D7_0809100 (an upsB type *var* gene; highlighted in bold), which has been shown to be translated in *P. falciparum* NF54 sporozoites in (*79*) but not in (*25*), is detected in female gametocyte and ookinete cells. Additionally, PF3D7_1200600 (*var2csa*; highlighted in bold) which has been primarily associated with pregnancy-associated malaria and has recently been shown to be expressed in male gametocyte bulk transcriptomic data (*6, 80*) was here detected in a single male gametocyte. The majority of cells where we detected sense *var* transcripts displayed mutually exclusive expression (expressing a single *var* gene). However, we detected simultaneous expression of two *var* genes in four cells (female gametocytes and sporozoite), but in these cases, one or both *vars* were expressed at very low levels (fewer than 5 sense reads detected).

**Figure S10.**
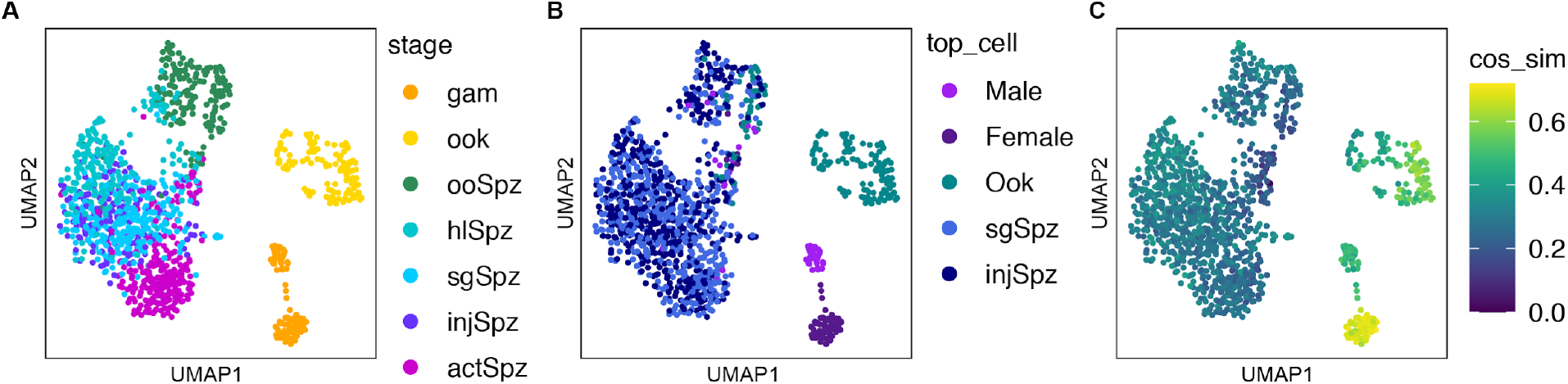
scmap of *P. falciparum* transcriptomes to *P. berghei* dataset. Scmap (*14*) was used to compare our data with equivalent stages from the *P. berghei* data set (*8*). A reference index for *P. berghei* was built based on one-to-one orthologs and the *P. falciparum* data was mapped to it to match each cell to a *P. berghei* cell. **(A-C)** UMAP of *P. falciparum* cells coloured according to collection stage **(A)**, stage of the matched *P. berghei* cell **(B),** and cosine similarity metric for each cell **(C)**. In general, cells matched to a cell from a similar stage in the *P. berghei* dataset. Female gametocytes and ookinetes had a higher cosine similarity on average compared to sporozoites and male gametocytes, which also tend to have fewer genes per cell.

**Figure S11.**
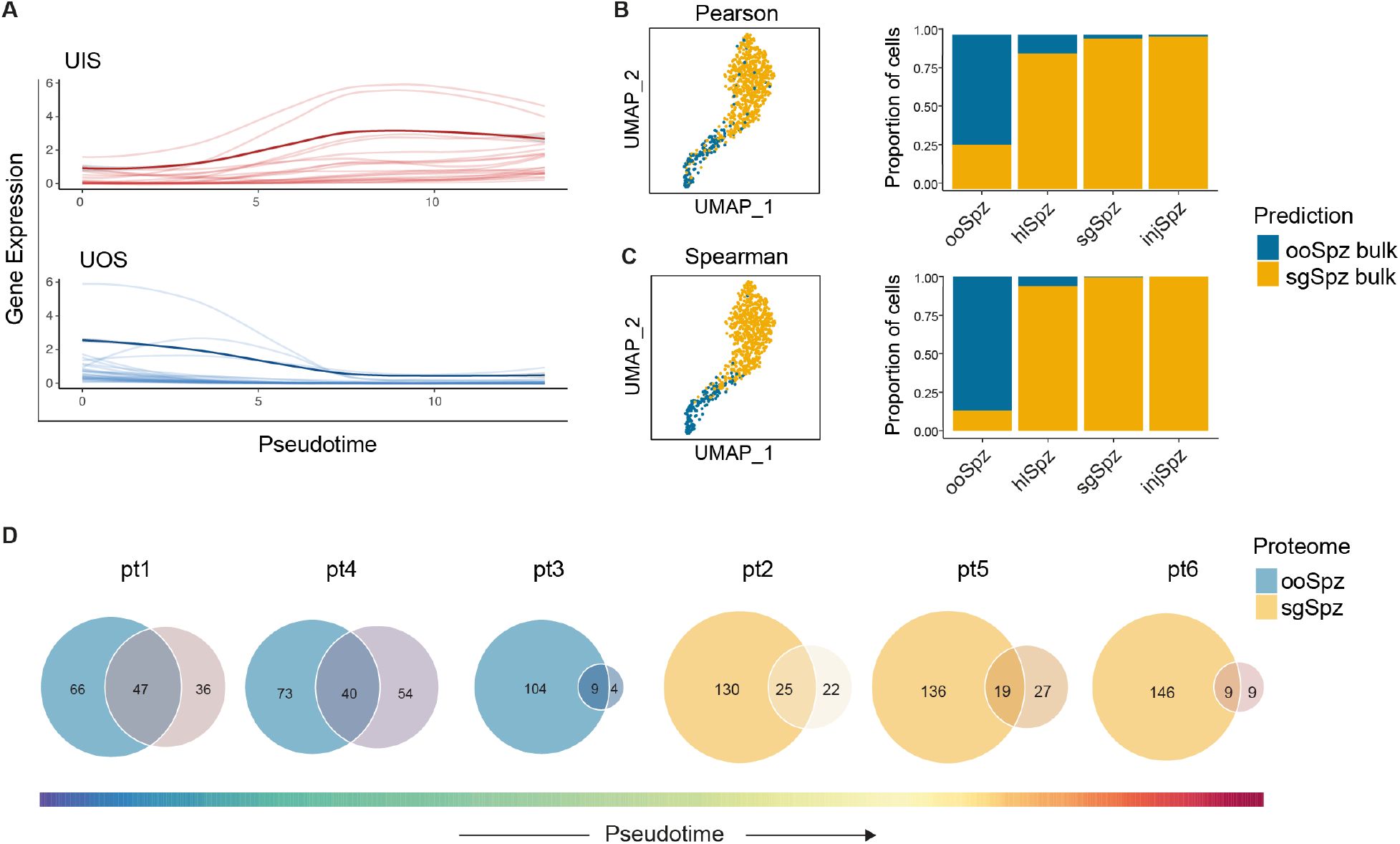
Comparison of the sporozoite transcriptional programme between single-cell and bulk data sets. **(A)** Expression dynamics of 21 UIS (top) and 29 UOS (bottom) genes identified by Lindner *et al. (25*) that show DE over pseudotime (q-val <0.0001). Individual genes are represented by light coloured lines and the average expression of each gene set is shown in darker colours. (**B, C**) Predicted sporozoite stage based on Pearson (**B**) and Spearman (**C**) correlations with available bulk data. The left and right panels show UMAP representations of single-cell transcriptomes coloured by predicted stage and the quantitative distribution of cells according to their predicted stage, respectively. Single-cell transcriptomes matched to either oocyst or salivary gland bulk data as expected. (**D**) Correspondence between mRNA and protein expression for DE genes over pseudotime. Proteins where the gene showed differential expression over pseudotime were selected from the two proteome datasets (ooSpz and sgSpz) from (*25*) and compared to each pseudotime gene cluster (pt 1-6) from the same stage. The Venn diagrams show the intersection of each cluster of DE genes (pt 1-6) with ooSpz or sgSpz sub-proteomes (*25*). Sporozoite transcripts for which no protein products could be detected in Lindner *et al*. are likely to be translationally repressed.

**Figure S12.**
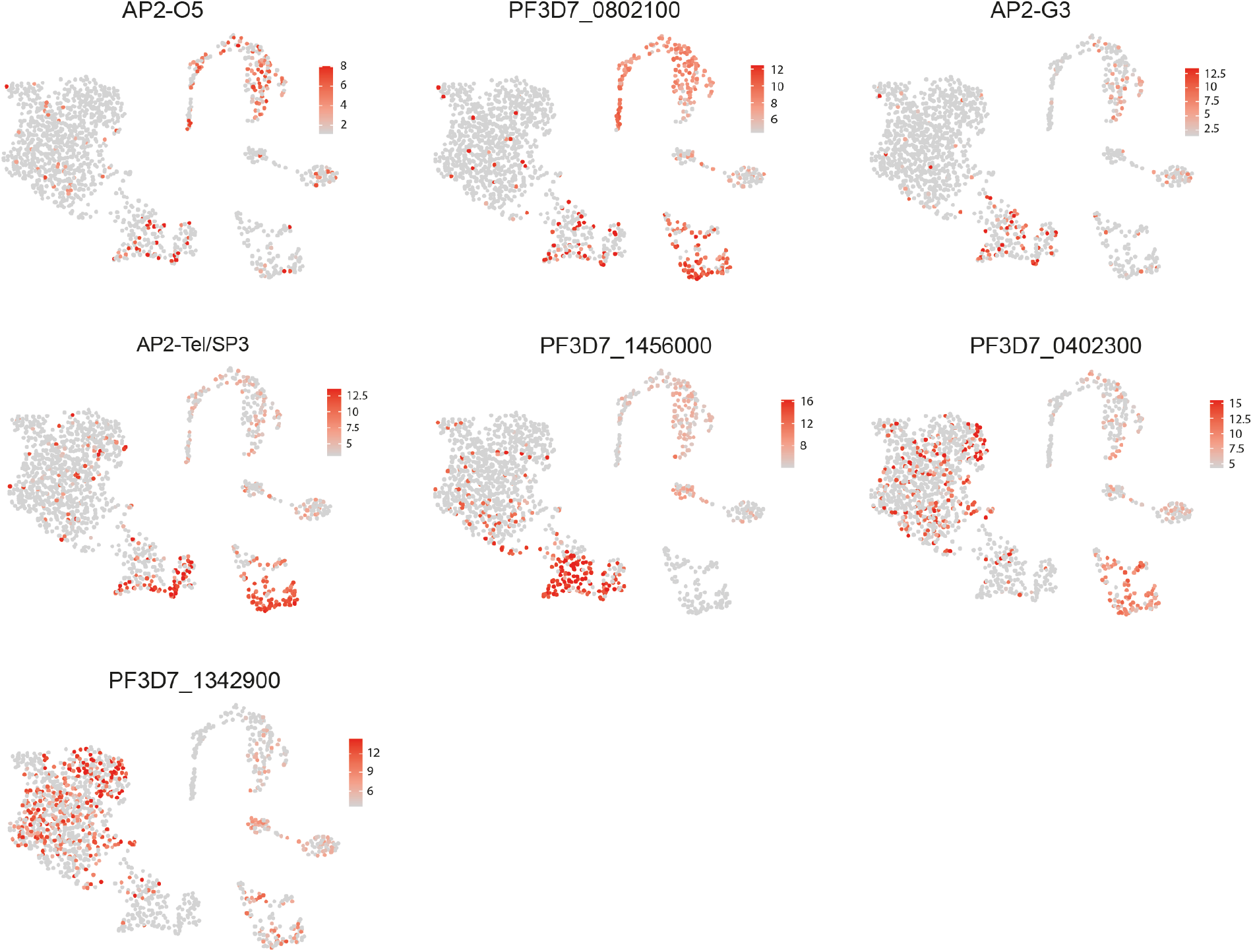
ApiAP2 transcription factors associated with sporozoite development in the mosquito. UMAP representations of the parasite life cycle with AP2 transcription factors DE along sporozoite pseudotime highlighted in red. Expression levels are colour-coded according to the scale next to each plot. Only AP2Tel/Sp3 (PF3D7_0622900) has been previously implicated in sporozoite development in *P. berghei* (*33*). AP2-G3 (PF3D7_1317200) and AP2-O5 (PF3D7_1449500) have known roles in sexual development and ookinete motility, respectively (*34*).

**Table S1.**
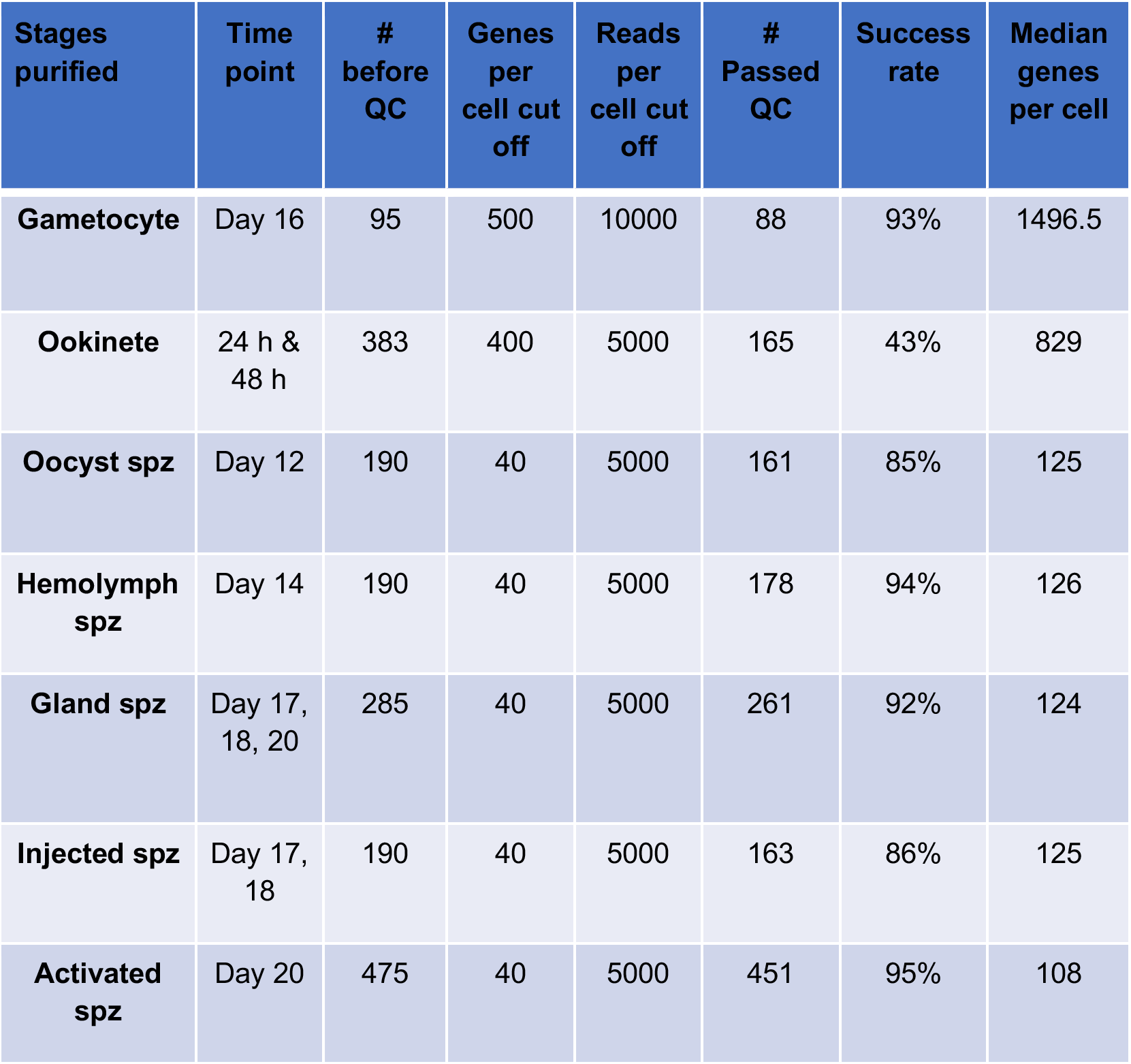
Quality control of single-cell transcriptomes by isolation method. Gametocyte time point is represented as days post gametocyte induction, all other time points are hours or days post infectious feed.

## Acknowledgements

The authors wish to acknowledge the support from the FACS facility from Imperial College London, namely Jane Srivastava and Radhika Patel, for single-cell sorting and the core sequencing pipelines at the Wellcome Sanger Institute. We thank Mark Tunnicliff for maintenance and provision of *Anopheles stephensi* mosquitoes and Alisje Churchyard, Sabrina Yahiya, and Catherine Ducker for assistance with experiments or figures.

## Author contributions

E.R., V.M.H., F.D. and K.W designed and performed all experiments and analysed the data. J.C., C.A.B, A.J.R. and M.D. helped with computational analysis of data and figure preparation. S.K.D. and V.M.H. built the interactive web tool. J.B. and M.K.N.L. sourced funding, provided scientific direction and helped with experimental design, analysis of data, and manuscript preparation. E.R and V.M.H. wrote the manuscript with contributions from other authors.

## Data Availability

Raw sequence data are available from the European Nucleotide Archive (accession number ERP124136). Expression matrices and supporting files are available on Github at https://github.com/vhowick/pf_moz_stage_atlas and data are searchable via the MCA website:www.malariacellatlas.org.

